# The pivotal role of Drgx in survival, wiring and identity of T4/T5 neurons

**DOI:** 10.1101/2024.03.12.584653

**Authors:** Laura Gizler, Katharina Schneider, Sarah Steigleder, Simon Benmaamar, Stephan Schneuwly, Mathias Rass

## Abstract

The development of *Drosophila melanogaster’s* T4/T5 motion-sensing neurons has been extensively studied. Despite identifying many genes, important developmental steps remain unknown. This study investigates the Paired-like homeobox transcription factor Dorsal root ganglia homeobox (Drgx) in T4/T5 neuron development. *Drgx* expression initiates in T4/T5 neuroblasts and persists in mature neurons. Knockdowns using T4/T5-specific drivers yield distinct phenotypes in the optic lobe, including dendrite mistargeting and reduced lobular plate size due to apoptosis in early knockdowns. Late knockdowns exhibit only extensive mistargeting. Therefore, Drgx plays a dual role, initially inhibiting apoptosis and later on establishing T4/T5 neuron identity and circuit formation. Targeted DamID (TaDa) and RNA-sequencing (RNA-seq) identify Drgx target genes involved in apoptosis and neuron projection development. Therefore, Drgx orchestrates vital stages in T4/T5 neuron development, influencing survival, identity, and circuitry and connects to the previously identified transcription factor *Sox102F* as a target of Drgx.

**Summary statement:** The Paired-like homeobox transcription factor Drgx is essential for the correct establishment of the identity of a specific type of motion vision neurons in *Drosophila*.

## Introduction

The formation of the correct circuitry during the development of the nervous system is not only essential for the proper function of the nervous system but also important for the survival of the neuron itself (Levi-Montalcini, 1987; Oppenheimer, 1991). The finding of the correct neuronal partner depends on the expression of specific cell surface molecules that identify the different neurons (Carrillo et al., 2015; Özel et al., 2021; Tan et al., 2015; Xu et al., 2022). The expression of these molecules, in turn, is regulated by the developmental gene cascade defining the identity of the neuron (Konstantinides et al., 2022; Kurmangaliyev et al., 2020). One of the most powerful tools to study the formation of neuronal circuits is the visual system of *Drosophila melanogaster* due to its clear structure and well-studied synaptic connections (Courgeon and Desplan, 2019; Fischbach and Dittrich, 1989; Ngo et al., 2017; Takemura et al., 2008). Especially the formation of the motion vision pathway with its defined dendritic and synaptic interactions has evolved into a popular system to study the mechanisms leading to proper connectivity (Contreras et al., 2018; Oliva et al., 2014; Schilling et al., 2019; Silies et al., 2014).

The motion vision pathway is comprised, among other things, of T4/T5 neurons, which have their cell bodies adjacent to the lobula plate. The T4 and T5 neurons are defined by their different contrast polarity response, whereas the T4 neurons respond to brightness increment (ON-pathway) and the T5 neurons respond to brightness decrement (OFF-pathway) (Borst, 2014; Maisak et al., 2013). The T4 neurons send their dendritic connections into the medulla layer M10 and the T5 dendrites are found in the lobula layer Lo1 (Fischbach and Dittrich, 1989). For the further computation of the optomotor signal, the T4/T5 neurons signal back into the four lobula plate layers. Thereby each lobula plate layer is only innervated by one of the four T4/T5 neuronal subtype (a-b). Each subtype is responsible for signaling movement information in one of the four cardinal directions (front-to-back, back-to-front, upwards, and downwards) (Maisak et al., 2013).

The development of T4/T5 neurons has been studied intensively in the last years (Apitz and Salecker, 2015; Apitz and Salecker, 2018; Mora et al., 2018; Pinto-Teixeira et al., 2018). They originate from the inner proliferation center (IPC) which develops together with the outer proliferation center (OPC) during early L1 larval stages (Hofbauer and Campos-Ortega, 1990; Nassif et al., 2003). Both are built up by neuroepithelial cells that develop into neuroblasts (Egger et al., 2007; Green et al., 1993). The OPC derived neuroblasts give rise to neurons of the lamina and medulla (Apitz and Salecker, 2014; Yasugi et al., 2008). Neuroblasts of the IPC are the base for lobula neurons, the so-called C/T neurons, which are comprised of the T2a, T2, T3, C2 and C3 neurons, and the T4/T5 neurons (Hofbauer and Campos-Ortega, 1990; Mora et al., 2018; Oliva et al., 2014).

The IPC is further divided into four different subpopulations, the proximal-IPC (p-IPC), which is interconnected with the surface-IPC (s-IPC), the distal-IPC (d-IPC) and migratory streams connecting the p-IPC and the d-IPC (Apitz and Salecker, 2015). In a first step during the T4/T5 neuronal development, the s-IPC expresses Wingless (Wg) which is responsible for the expression of *decapentaplegic* (*dpp*) in the ventral and dorsal part of the p-IPC. Dpp then represses the default expression of *brinker* (*brk*) in these parts of the p-IPC creating two different cell populations. It also leads to the transformation of the neuroepithelial cells into the migrating progenitor cells via an epithelial-mesenchymal transition (EMT)-like process (Apitz and Salecker, 2015; Apitz and Salecker, 2018). From both p-IPC populations, cells migrate into the d-IPC and start to express *Dichaete* (*D*). After reaching the d-IPC the cells mature into neuroblasts and produce ganglion mother cells (GMCs) via asymmetric cell division (Apitz and Salecker, 2015).

During the development in the d-IPC, two competence windows can be defined that are created by a temporal relay. In the first window, cells in the lower d-IPC, that are located closely to the migratory streams, express *D* and *asense* (*ase*) generating Prospero positive GMCs, which produce the different C/T neurons (Apitz and Salecker, 2015). In the second competence window, the T4/T5 neurons are created from cells further away from the migratory streams. These neuroblasts are Tailless (Tll) positive and express the two transcription factors *dachshund* (*dac*) and *atonal* (*ato*). With the next division two ganglion mother cells (GMC) are produced, which divide once more to create two post mitotic T4/T5 neurons (Apitz and Salecker, 2015; Oliva et al., 2014; Pinto-Teixeira et al., 2018).

The eight different types of T4/T5 neurons are created in three steps during this development. In the first step the fate between T4/T5a,b and T4/T5c,d is induced. This decision depends on the Dpp expression in the p-IPC, which turns on the expression of *optomotor blind* (*omb*), not only in the ventral and dorsal p-IPC, but also in their progenitor cells in the d-IPC. In the T4/T5 GMCs, Omb suppresses the expression of *dac* leading to the formation of a T4/T5c,d fate. The GMC, which originates from p-IPC cells that do not express *dpp*, keep the expression of *dac* and obtain a T4/T5a,b fate (Apitz and Salecker, 2018). The next two decisions depend on the different Notch (N) states of the cells. First the inherited N state of the GMC decides the fate between the different subtypes (a or b, c or d). The GMCs that inherit the active N-pathway produce either T4a/T5a or T4d/T5d neurons. Respectively, the GMC in which the N-pathway is turned off due to the inheritance of the N inhibitor Numb produces T4b/T5b or T4c/T5c neurons. In the last step, the fate between T4 and T5 neurons is decided. This also depends on the N status of the cell. T4 neurons are produced in the absence of N and T5 neurons depend on the presence of N (Pinto-Teixeira et al., 2018).

In addition to the genes mentioned above, many other genes have been identified in recent years that are necessary for the formation of the T4/T5 neuron identity, e.g., *Lim1*, *acj6*, *SoxN* and *Sox102F* (Kurmangaliyev et al., 2019). In particular, the effects of *Sox102F* on the development of T4/T5 neurons have been intensively studied by different groups (Contreras et al., 2018; Schilling et al., 2019). In these studies, they could show that the mutation and the knockdown of *Sox102F* leads to a wrong connectivity of the T4/T5 neurons in the medulla and the lobula. Here, their dendrites overgrow their desired stopping layer M10 and Lo1 (Schilling et al., 2019). Additionally, in *Sox102F* knockdowns, the lobula plate size is reduced and ectopic bridges are found between the lobula and the lobula plate (Contreras et al., 2018). These results led to the conclusion that Sox102F controls the correct circuit formation and terminal differentiation of the T4/T5 neurons.

In this paper we investigate the function of an additional transcription factor, Dorsal root ganglia homeobox (Drgx), which has been identified to be expressed in T4/T5 neurons in different single cell RNA-sequencing (RNA-seq) data sets (Davie et al., 2018; Kurmangaliyev et al., 2019) and which potentially represents an important new link for the development of T4/T5 neurons. We confirm the expression of *Drgx* in T4/T5 neurons, thereby showing that *Drgx* is already expressed in T4/T5 neuroblasts. In addition, the knockdown of *Drgx* in T4/T5 neurons leads to the loss of the optomotor response in adult flies and histological analysis shows severe structural changes in the adult optic lobe. A T4/T5 neuron specific Targeted DamID (TaDa), as well as a T4/T5 neuron specific RNA-sequencing (RNA-seq), revealed many potential Drgx regulated genes. Interestingly, we were able to identify a Drgx binding side in the *Sox102F* locus and a significant down regulation of *Sox102F,* which fits very well with the common phenotypes of *Drgx* and *Sox102F* knockdowns. In the end we demonstrate that Drgx is positioned upstream of *Sox 102F* and regulates its expression. In summary, our findings support the conclusion that Drgx plays a significant role in determining the fate of T4/T5 neurons and that it is indispensable during the initial neuroblast stages.

## Results

### Drgx expression analysis reveals expression in T4/T5 neurons and perineurial glial cells in the larval and adult brain

To further understand the molecular and developmental cascade involved in establishing the T4/T5 neuronal circuits we screened the single cell RNA-seq data sets from Davie et al. (2018) and Kurmangaliyev et al. (2019) to identify genes that are highly specific for T4/T5 neurons and not previously associated with their development. One of these genes we discovered is *Drgx*. *Drgx* is located on the left arm of chromosome 2, and two splice forms have been identified, differing solely in the 3’ UTR. To investigate *Drgx* expression and function, we used two MiMIC lines, Mi11472 and Mi06689, in addition to the UAS-Drgx-IR knockdown line (dsRNA-HMC04912) (Fig. 1A). Mi11472 is located in the first intron and results in GFP expression and a premature termination of *Drgx* transcription. We performed recombination-mediated cassette exchange to eliminate the MiMIC cassette and replaced it with a *GAL4* gene. This *GAL4* construct will be regulated by the *Drgx* promoter and facilitate the overexpression of UAS-transgenes, such as fluorophores, to investigate the expression profile of *Drgx*. To confirm the GAL4 line’s expression pattern, we replaced the MiMIC cassette of Mi06689, located in intron 6 of *Drgx*, with GFP. This resulted in the expression of a Drgx::GFP fusion protein (Fig. 1A).

**Fig. 1:**
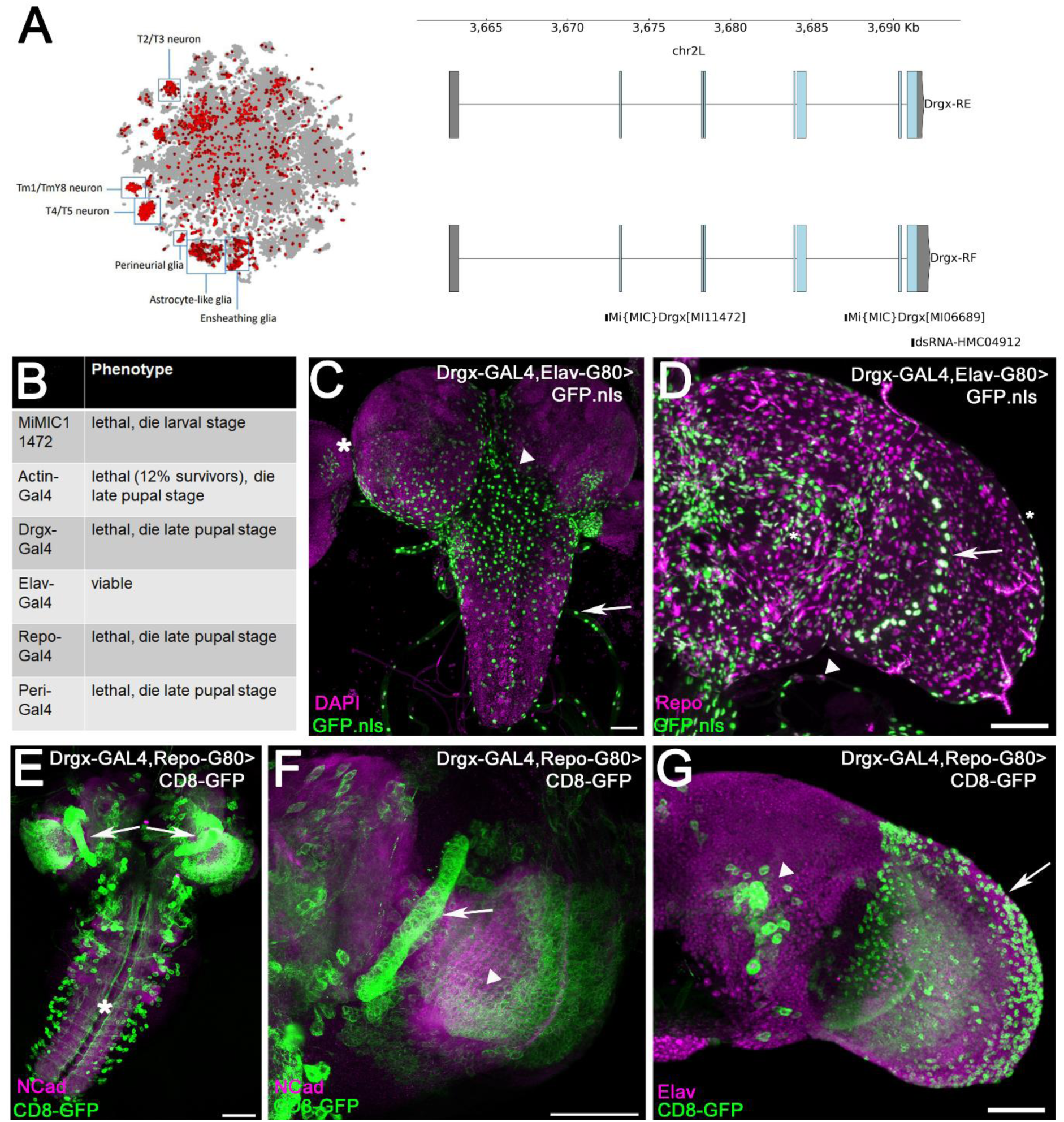
Drgx expression is found in neurons and glial cells of the larval and adult brain. (A) tSNE plot from SCope showing *Drgx* expression in the adult *Drosophila* brain, based on single-cell RNA-seq from Davie et al. (2018). Based on this dataset, *Drgx* is expressed in T4/T5, Tm1/TmY8, and T2/T3 neurons, as well as ensheathing-, astrocyte-like-, and perineurial-glial cells. The gene *Drgx* is located on chromosome 2L and produces two transcripts that differ solely in the 3’ UTR. The two main MiMIC lines are Mi11472 and Mi06689, whereby Mi11472 is integrated in the 5’ UTR intron in orientation of the reading frame and Mi06689 is situated in intron 5, opposite the reading frame. The RNAi line (HMC04912) targets a sequence in exon 7. (B) First knockdown experiments show lethality with different general driver lines and with Drgx-GAL4. (C) Restriction of Drgx-GAL4 expression to glial cells with Elav-G80 shows expression throughout the brain (arrowhead), eye discs (asterisk) and nerve fibers (Arrow). (D) In the adult optic lobe, expression is observed in the outer most glial cells layer of the brain (asterisk) and the nerve fibers (arrowheads) and in the giant glial cells become visible (arrow). (E) The neuronal pattern of Drgx generated with Drgx-GAL4/UAS-CD8-GFP; Repo-G80 can be observed in the larval optic lobes (arrow) and ventral nerve cord (asterisk). (F) In the developing larval optic lobe, Drgx expression is present in both the dIPC (arrow) and the developing T4/T5 neurons (arrowhead). (G) In the adult brain, a Drgx-positive cell cluster is present in the central brain (arrowhead) in addition to its expression in the optic lobe. Scale bars: 50 µm.

Late-stage larval lethality was observed in homozygous Mi11472 larvae (Fig. 1B). To confirm that the lethality was due to early termination of *Drgx* transcription, rather than a potential background genetic effect of this line, we performed an actin-GAL4 knockdown of *Drgx* in all cells. This resulted in late pupal lethality with only 12% of the flies surviving to adulthood. Similarly, late pupal death was observed in homozygous Drgx-GAL4 flies. To determine the specific cells responsible for the pupal lethality of the *Drgx* knockdown, we used different GAL4 lines to express the Drgx-IR knockdown in different cell types. Pupal lethality was not observed when *Drgx* was knocked down in all neurons; however, *Drgx* knockdown was lethal when expressed in all glial cells. We subsequently knocked down *Drgx* in all glial subtypes and found that late pupal lethality was caused by the knockdown of *Drgx* in perineurial glial cells (Fig. 1B). To visualize Drgx expression in glial cells, we used the Drgx-GAL4, Elav-G80 line to express nuclear GFP (GFP.nls), preventing GFP.nls expression in neurons. In larvae, *Drgx* is expressed only in a specific subset of glial cells located on the surface of the brain, in the eye imaginal disc, and in the ganglia. Based on this expression pattern *Drgx* is primarily expressed in perineurial glial cells (Fig. 1C). In adult flies, *Drgx* is expressed in the perineurial glia as well as in the inner giant chiasm glial cells (Fig. 1D). To visualize neuronal *Drgx* expression, we used a Drgx-GAL4, Repo-G80 line to express membrane-anchored UAS-CD8-GFP, which suppresses GFP expression in glial cells. In larvae, UAS-CD8-GFP expression was found in single neurons in the hemineuromeres of the ventral nerve cord (Fig. 1E) and broadly in the developing optical lobes (Fig. 1F). In adults, we observed UAS-CD8-GFP expression in antennal lobe neurons and especially in the optical lobes (Fig. 1G). Based on the positioning of these neurons in both larval and adult optic lobes, it is plausible to suggest that these cells may be transmedullary Tm neurons of the medulla and T4/T5 neurons of the lobula plate. The Drgx-GAL4 gene expression detected with the GFP variants validates our original findings, which were subsequently confirmed with the Drgx::GFP line.

## *Drgx* is expressed in d-IPC Neuroblasts and all descending T4/T5 neurons

To characterize the neurons expressing Drgx in the developing optic lobe, we used larval brains expressing Drgx::GFP and stained them with anti-Dac, a marker for young T4/T5 neurons in L3 larvae (Apitz and Salecker, 2018). Drgx::GFP expression was observed to overlap with Dac expression in T4/T5 neuroblasts and neurons, but not in Dac positive medulla neurons (Fig. 2A). To confirm Drgx expression in T4/T5 progenitors, we stained brains of Drgx::GFP expressing larvae with anti-pHH3, a marker for mitosis, and again found colocalization in the dIPC area (Fig. 2B). Colocalization experiments using Drgx::GFP and Abnormal Chemosensory Jump 6 (Acj6) show that Drgx is expressed not only in T4/T5 neurons, but also in developmentally related C/T neurons (Fig. 2C).

**Fig 2:**
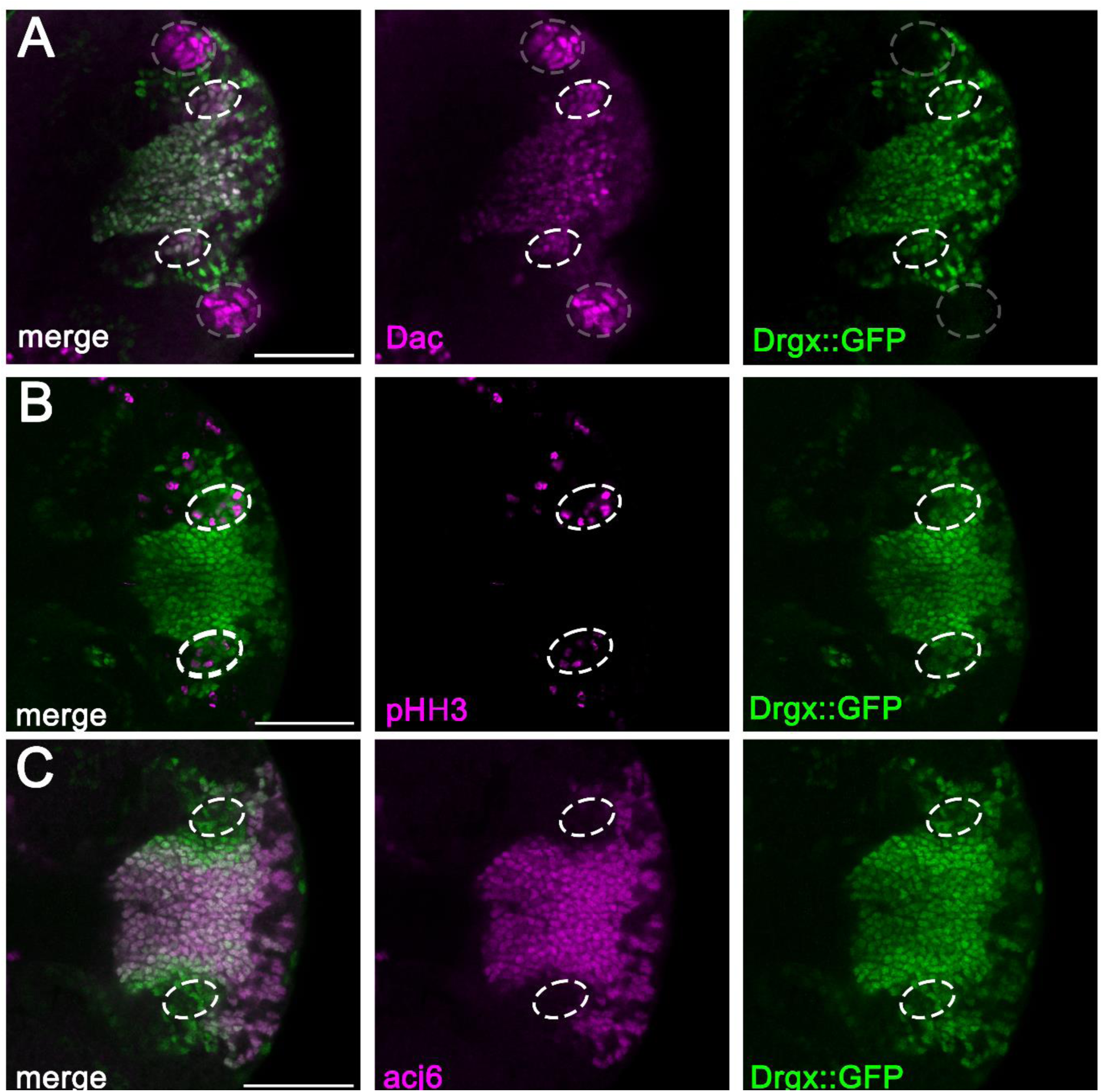
*Drgx* is expressed in dividing T4/T5 neuroblasts of the dIPC. (A) A fraction of the cells stained with anti-Dac in the developing third instar larval optic lobe overlap with Drgx::GFP expression (white circles), indicating *Drgx* expression in T4/T5 neuroblasts in the dIPC. On the right side of the optic lobe, there are Drgx::GFP positive cells that do not express Dac, which could potentially represent C/T neurons. In addition, Dac staining identified cells that are Dac positive but Drgx negative, which belong to the lamina precursor cells in the OPC (gray circle). (B) Drgx::GFP expression overlaps with pHH3 in the dIPC, indicating that Drgx is expressed in dividing cells, which are potential T4/T5 neuroblasts. (C) Near the developing, Acj6-positive T4/T5 neurons in the larval optic lobe, Acj6-negative cells can be observed expressing Drgx::GFP. These cells correspond to the developmentally related C/T neurons. Scale bars: 40 µm.

### Knockdown of *Drgx* in T4/T5 neurons results in loss of optomotor response

Next, we examined whether the Paired-like homeobox transcription factor Drgx plays a crucial role in the development of T4/T5 neurons and, subsequently, their function in adult flies. These neurons are responsible for detecting motion in *Drosophila* (Borst, 2014; Maisak et al., 2013). Since our previous findings indicated that *Drgx* knockdown in all neurons is not lethal, we conducted tests on the optomotor behavior of flies and demonstrated that they lacked a response (Fig. 3A). The optomotor response can be quantified by the optomotor magnitude, which reflects the strength of the response, and the optomotor slope, which indicates its speed. In contrast to control flies, both parameters were significantly diminished in pan-neuronal *Drgx* knockdown (Fig. 3A). We performed the experiment by silencing *Drgx* expression in all developing T4/T5 neurons using the R40E11-GAL4 driver line to rule out the possibility of other *Drgx*-expressing neurons besides T4/T5 being accountable for this phenotype. This driver line has been previously confirmed to be highly effective in T4/T5-specific optomotor experiments (Schilling et al., 2019). Our results for the T4/T5 neuron-specific knockdown of *Drgx* demonstrate a significant reduction in optomotor magnitude and optomotor slope, resulting in the complete absence of the optomotor response (Fig. 3B). Thus, our findings confirm that *Drgx* expression in T4/T5 neurons is critical for accurate motion vision. To determine whether *Drgx* knockdown flies retained any visual function, we performed the Buridan’s paradigm analysis. We placed flies with clipped wings in a white illuminated arena. The only visual landmarks are two black stripes that face each other, which the flies are unable to reach because the arena is surrounded by water (Colomb et al., 2012). In flies used in this experiment, *Drgx* was knocked out during development in all neurons using Elav-GAL4 or in developing T4/T5 neurons using R40E11-GAL4. Despite the knockdown, the experimental flies walked between the stripes similar to the controls, indicating that their vision remained unaffected (Fig. S2).

**Fig. 3:**
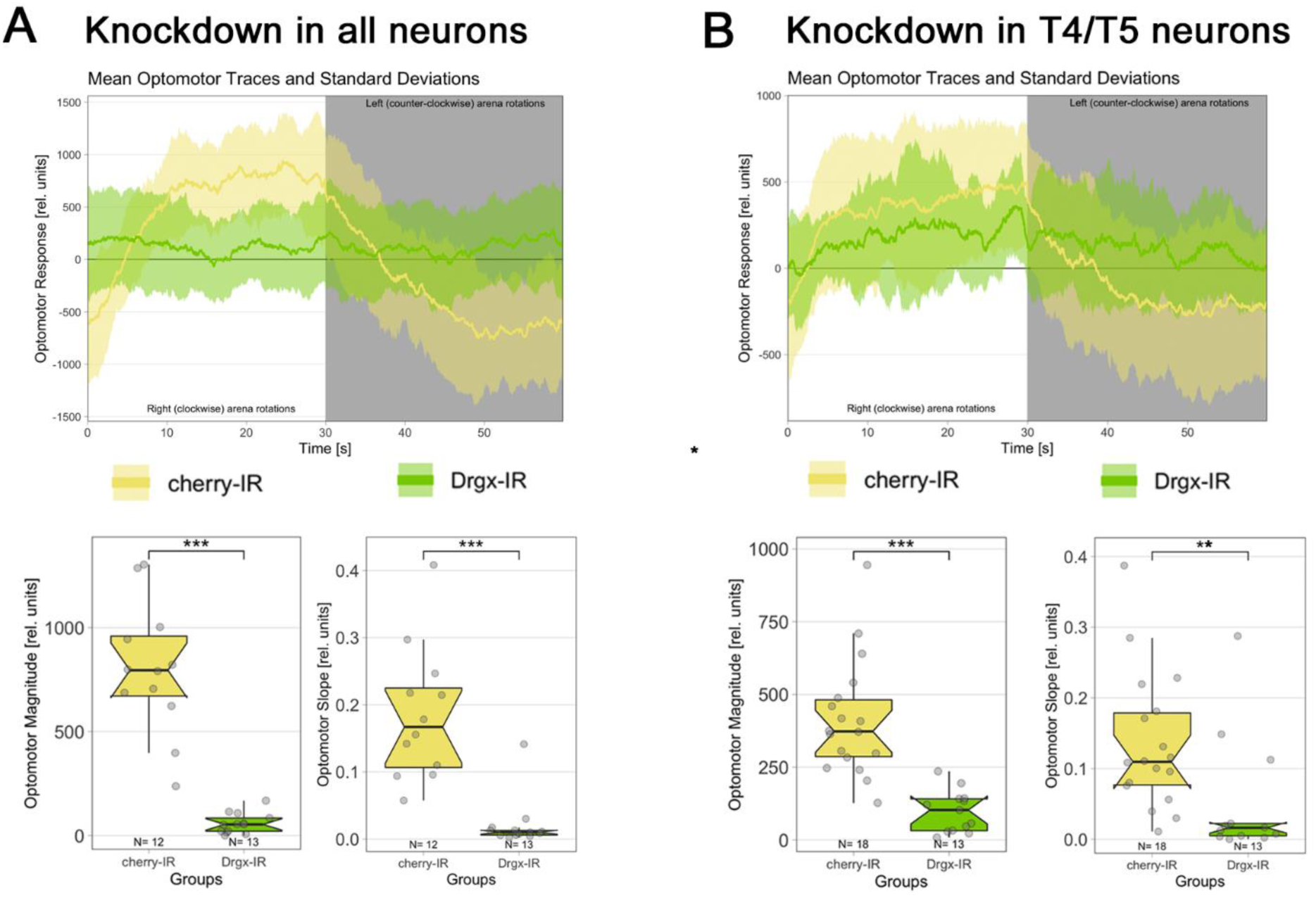
Pan-neuronal and T4/T5 specific *Drgx* knockdown abolishes the optomotor response. (A) The pan-neuronal knockdown of *Drgx* using Elav-GAL4 reveals absence of a double sigmoidal curve in the mean optomotor trace, as seen in the control knockdown of *cherry*. The optomotor magnitude is significantly lower in flies with *Drgx* knockdown (n=12, p-value=3.85e-07). Additionally, the optomotor slope is significantly smaller when *Drgx* knockdown is expressed (n=13, p-value=4.62e-06) compared to the control optomotor slope. (B) In the T4/T5 specific optomotor experiment, control flies (UAS-cherry, R40E11-GAL4) exhibited the expected turning behavior, as indicated by the mean optomotor traces with standard deviation. Knockdown flies did not exhibit this behavior, showing almost no turning in response to changes in direction. The calculated optomotor magnitude of *Drgx* knockdown flies is significantly lower than that of the control group (n=18, p-value=2.91e-07). A significant reduction in optomotor slope was also observed in the *Drgx* knockdown flies (n=13, p-value=0.00271). For statistical analysis Mann-Whitney U-tests were performed.

### Early knockdown of *Drgx* in T4/T5 neurons and their progenitors leads to reduction of adult lobula plate size

Given that the optomotor response was completely lost in flies with *Drgx* knockdown in developing T4/T5 neurons, we were intrigued to see how the knockdown affected the morphology of T4/T5 neurons and the lobula plate, where the T4/T5 neurons terminate. In order to investigate this, we used three different driver lines. Two are already expressed in T4/T5 precursors and their descending neurons, called R9B10-GAL4 (Apitz and Salecker, 2018) and R23G12-GAL4 (Kurmangaliyev et al., 2019). The third, called R39H12-GAL4 (Schilling et al., 2019), is expressed in older maturing T4/T5 but not in their neuroblasts or GMCs. Adult brains were stained using anti-Bruchpilot (Nc82) to visualize active synaptic zones and thus the neuropils of the optic lobe, and membrane-bound GAL4-driven UAS-CD8-GFP was used to visualize the morphology of T4/T5 neurons. In control flies (R9B10-GAL4; UAS-cherry-IR and R23G12-GAL4; UAS-cherry-IR, R39H12-GAL4; UAS-cherry-IR), the three optic neuropils, including the lobula, lobula plate, and medulla, were visible (Fig. 4A, Fig. 4C, Fig. 4E). UAS-CD8-GFP expression revealed the dendritic connections of T4/T5 neurons in lobula layer 1 and medulla layer 10. Since the R9B10-GAL4 resulted in very weak GFP expression in adults (Fig. 4A) and the GFP signal was even further reduced in experimental flies to almost background levels (Fig. 4B), the second early driver line R23G12-GAL4 (Fig. 4C, Fig. 4D) was used, which showed stronger GFP expression in adults. In *Drgx* knockdown with either of the two early lines, the lobula and medulla were similar in structure to controls, however, the size of the lobula plate appeared to be strongly reduced, its structure was partially disrupted, and part of the lobula plate was fused to the lobula (Fig. 4B, Fig. 4D). These findings are consistent with cell death. In addition, the dendritic connections in the lobula and medulla are mistargeted to deeper layers of these neuropils (Fig. 4D). When *Drgx* knockdown was induced later during T4/T5 development using R39H12-GAL4, the size and structure of the lobula plate was not affected, but as before, dendrites targeted not only lobula layer 1 and medulla layer 10, but also deeper layers (Fig. 4E, Fig. 4F). To see if the mistargeting effect was already visible at earlier stages of development, we examined the optic lobes of 24 hr old pupae and found that dendrites from knockdown flies were also partially mistargeted to deeper layers of the medulla and lobula in contrast to controls (Fig. S3). To evaluate the aforementioned reduction in the size of the lobula plate, we performed paraffin sections of adult fly heads. We measured the area of the lobula plate and divided it by the area of the medulla, which was used as a reference. Knockdown of *Drgx* with the panneuronal driver line Elav-GAL4 and the early T4/T5 expressing driver lines R9B10-GAL4 and R23G12-GAL4 resulted in a strong decrease in the size of the lobula plate (Fig. 4G), which was not observed with the late driver line R39H12-GAL4 (Fig. 4G). Since apoptosis of T4/T5 neurons is a prerequisite for lobula plate reduction and lobula plate-medulla fusion, we analyzed whether early *Drgx* knockdown leads to apoptosis of T4/T5 neurons. To visualize apoptosis, we used the apoptosis sensor GC3Ai (Schott et al., 2017). In developing T4/T5 neurons of controls, only a few single fluorescent cells can be observed (Fig. 4H, Fig. 4J and Fig. 4L), but after knockdown of *Drgx* with R9B10-GAL4 and R23G12-GAL4, the fluorescence signal is strongly increased (Fig. 4I, Fig. 4K), indicating that apoptotic events induced by *Drgx* knockdown have already occurred in larval T4/T5 neurons. When *Drgx* knockdown is shifted to a later stage of T4/T5 neuron development using the R39H12-GAL4 driver line (Fig. 4L), apoptotic events stay at control levels (Fig. 4M).

**Fig. 4:**
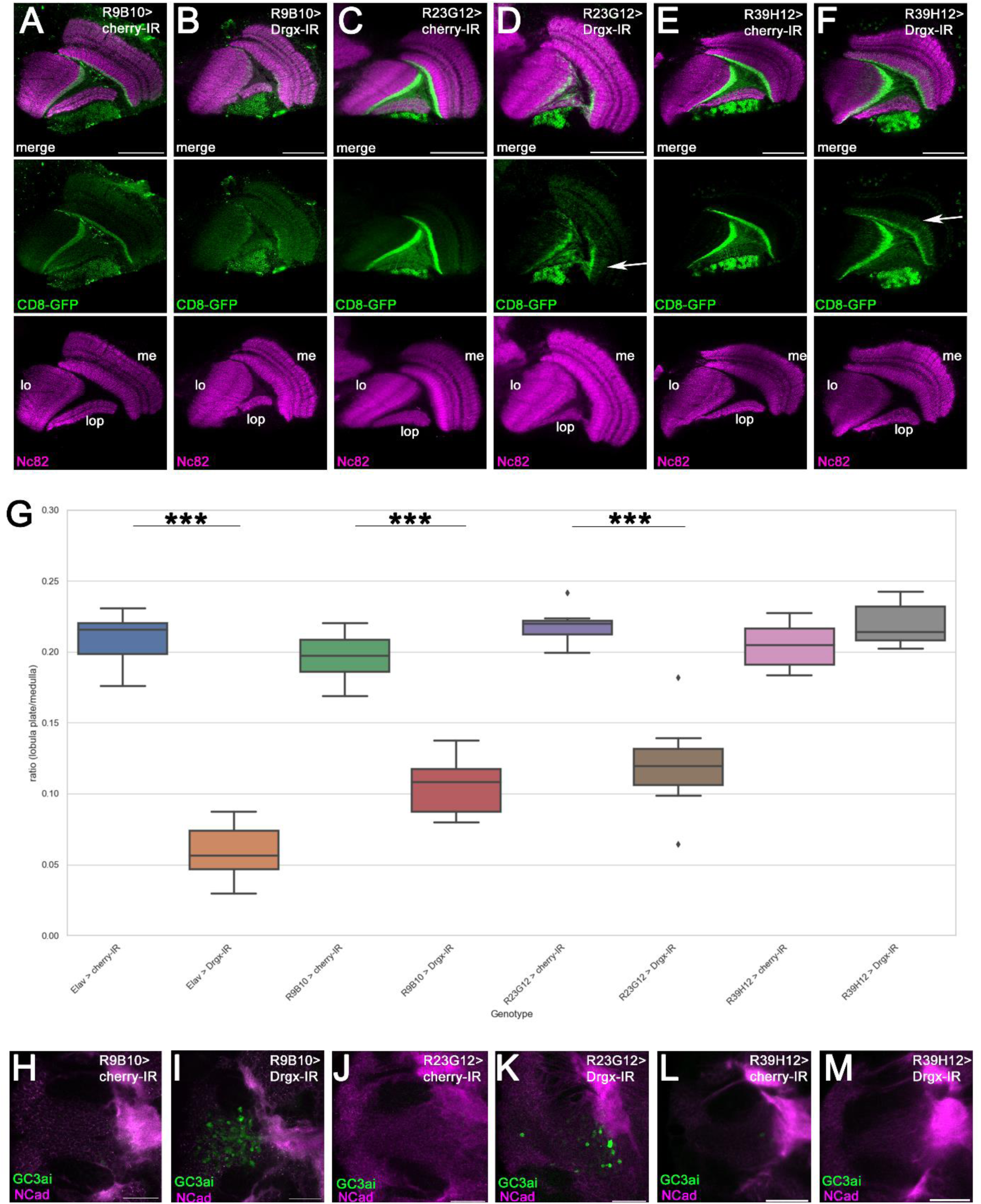
*Drgx* knockdown in T4/T5 neurons results in decreased lobula plate size due to apoptosis. (A, C, E) In the adult optic lobe, the neuropil structures of the lobula (lo), lobula plate (lop), and medulla (me) are visible in the control crosses with R9B10-GAL4, R23G12-GAL4, and R39H12-GAL4. (B, D) However, in the *Drgx* knockdown crosses with R9B10-GAL4 and R23G12-GAL4, the ordered structure of T4/T5 neurons breaks down. Specifically, dendrites overgrow their target layers in the lo and me (arrow), and the size of the lop reduces, and fuses with the lo. (F) In the *Drgx* knockdown with R39H12-GAL4, the size of the lop appears like that of the control group; however, the dendrites in the lo and me are still overgrowing their target layers (arrow). (G) To assess lop size reduction, the ratio of lop to me was calculated. The results demonstrated significant size reduction in *Drgx* knockdowns using Elav-G4 (n=15, p-value=3.3918e-06), R9B10-GAL4 (n=14-15, p-value=5.0970e-06) and R23G12-GAL4 (n=10, p-value=0.00018) compared to the control group, but not for R39H12-GAL4 (n=7-8, p-value=0.1520). Statistical analysis was performed using the Mann-Whitney U-Test. (H, J, L) Knockdown controls utilizing R9B10-GAL4, R23G12-GAL4, and R39H12-GAL4 did not exhibit any apoptosis in the developing optic lobe of the larval brain. (I, K) However, a noticeable increase in apoptosis was observed in the optic lobes of *Drgx* knockdowns utilizing the R9B10-GAL4 and R23G12-GAL4 driver lines. (M) Conversely, no increase in apoptosis was observed in the *Drgx* knockdown larval brains with R39H12-GAL4. Scale bars: 50 µm (A-F), 25 µm (H-M).

### TaDa Analysis revealed Drgx binding as a transcription factor to genes implicated in neuronal development

These results showed us that Drgx plays a significant role in the differentiation and development of T4/T5 neurons. Since Drgx is a homeobox transcription factor, our next approach was to investigate the binding pattern and binding sites of Drgx. To do this, we used the TaDa method (Southall et al., 2013) with a fusion protein of Drgx and the Dam methylase to identify binding sites of Drgx specifically in differentiating T4/T5 neurons. For this approach, we needed a GAL4 line that was highly specific for expression in T4/T5 neurons, and so we used the R23G12-GAL4 line, which shows very restricted expression only in developing T4/T5 neuroblasts and their descending neurons (Kurmangaliyev et al., 2019). As a control, we expressed only the Dam methylase with R23G12-GAL4 to detect background methylation. Averaged Drgx binding profiles across genetic regions showed enrichment at the transcription start site (TSS), as expected for a transcription factor, with enrichment decreasing along the gene body (Fig. 5A). After the transcription termination site (TTS), Drgx binding increases again (Fig. 5A). 2241 significant peaks were detected at 1% False Discovery Rate (FDR), which are very similar to the Drgx binding profiles. Thus, most of the peaks are found at the TSS, which is a strong indicator of transcriptional regulation (Fig. 5B). The majority of Drgx peaks are located 3kb upstream and downstream of the TSS (Fig. 5C). A total of 1579 genes were identified in the mapped peaks, and gene ontology (GO) enrichment analysis was conducted for biological processes and molecular functions. This analysis revealed that genes bound by Drgx are involved in biological processes such as development of neuron projections, morphogenesis of neuron projections, development of axons, and guidance of axons (Fig. 5D). When conducting GO enrichment analysis to cluster molecular functions, it was determined that Drgx binds to genes associated with RNA polymerase II regulation or transcription factor binding. This suggests that other transcriptional regulators are influenced by Drgx (Fig. 5E).

**Fig. 5:**
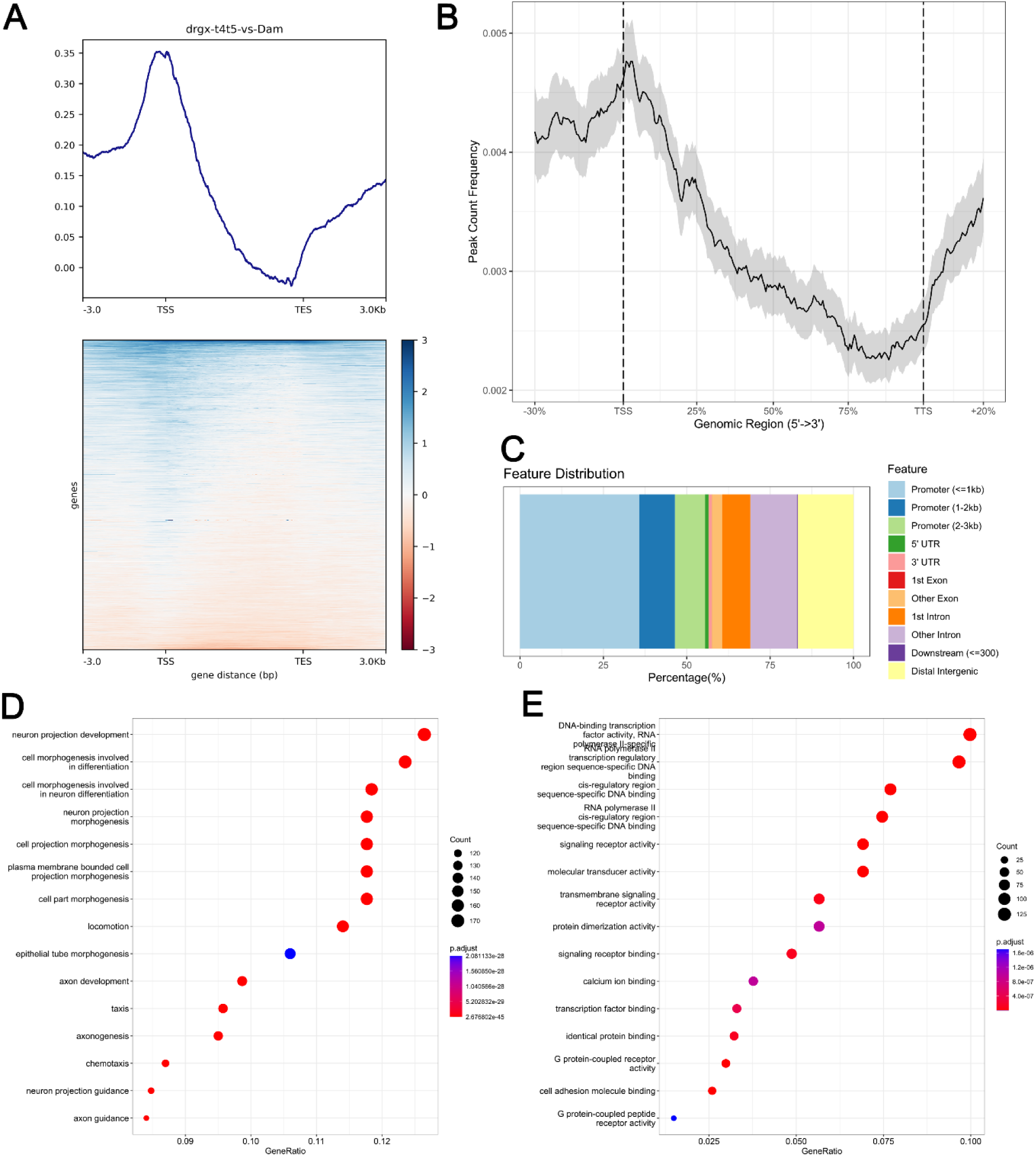
Drgx-Dam binding sites are close to the TSS of genes involved in neuron projection development and axon guidance. (A) Averaged Drgx-Dam/Dam signal across pseudogenic region and heatmap of DamID signal showing enriched binding of Drgx around TSS. (B) Drgx peak count frequency is increased around the TSS. (C) More than 50% of the Drgx peaks are observed 3 kb upstream or downstream of the promoter sites. (D) Gene ontology enrichment analysis revealed that Drgx-bound genes are involved in biological processes such as neuron differentiation, neuron projection development, axon guidance and development. The top 15 categories are shown. (E) Gene ontology enrichment analysis revealed that Drgx-bound genes have molecular functions such as RNA polymerase II regulation or transcription factor binding. The top 15 categories are shown.

### T4/T5 specific *Drgx* knockdown strongly affects the transcription of genes involved in neuron projection development and axon guidance

As DNA-binding of proteins to transcriptional start sites does not necessarily indicate whether a gene is up or downregulated, RNA-seq analysis of both control flies and experimental flies is necessary to determine how the transcriptome of T4/T5 neurons is affected after a *Drgx* knockdown. Third instar larvae brains with the corresponding genotype were dissected, and a single cell suspension was created. T4/T5 neurons were identified through UAS-CD8-GFP expression driven by R23G12-GAL4 and sorted using FACS. 901 genes had an adjusted P-value of less than 0.05 (Fig. 6A). The top 25 most significant DEGs included transcription factors (*scarecrow*, *Sox102F*, and *knot*) and genes involved in synapse formation (*DIP-delta, DIP-eta*, *DIP-eta*, *DIP-epsilon*, *Nlg2*, and *dpr9*) (Fig. 6A). Upregulated genes involved in apoptosis regulation were observed in T4/T5 neurons after a *Drgx* knockdown (Fig. 6B), which correlated with the reduction of lobula plate size. In addition, knockdown of *Drgx* led to T4/T5 neurons being mistarget. RNA sequencing data showed dysregulation in various protein families involved in pathfinding following a *Drgx* knockdown. Specifically, genes from the DPR, DIP, SIDE, BEAT, and NLG families were dysregulated (Fig. 6B), all of which play a role in synapse formation (Carrillo et al., 2015; Sink et al., 2001; Van Vactor et al., 1993; Xing et al., 2018). This observation is further supported by the GO enrichment analysis of biological processes. Similar to the genes bound by Drgx, the genes that show dysregulation after knockdown of *Drgx* with an adjusted p-value < 0.05, play a critical role in the development and morphogenesis of neuron projections (Fig. 6C). The GO enrichment analysis showed many terms that were also found in the TaDa GO analysis such as neuron projection development, neuron projection morphogenesis, neuron projection guidance and axon guidance (Fig. 6D). We histologically examined two early expressed axon pathfinding genes (*dpr11* and *Mp*) and a late expressed gene of unknown function (*CG14340*), which were found in both the TaDa and the RNA-seq data (Fig. S4-S5) and demonstrated the dependence of their expression on Drgx here as well.

**Fig. 6:**
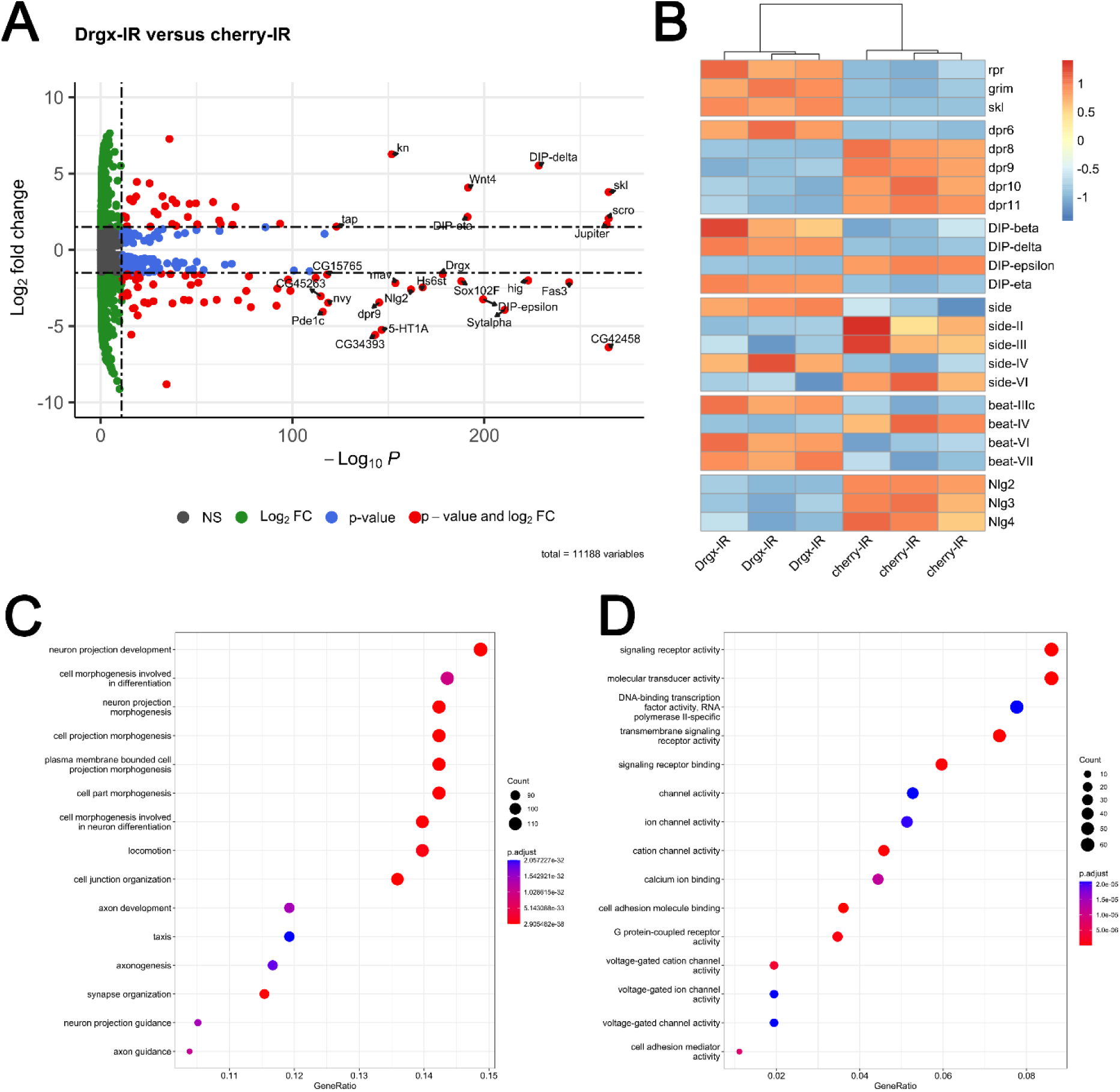
DEGs are involved in neuron projection development and axon guidance. (A) Volcano Plot showing the 25 most significant DEGs with a Log2FC ≥ 1.5 or a Log2FC ≤ *-*1.5 after *Drgx* knockdown in T4/T5 neurons. Red dots are genes with an adjusted p-value smaller than 10e-12 and a Log2FC ≥ 1.5 or a Log2FC ≤ *-*1.5. Blue dots only meet the p-value criteria. Green dots only meet the Log2FC criteria. Grey dots do not meet any criteria. (B) Heat map of DEGs belonging to different protein families involved in apoptosis (rpr, grim, skl) and axon guidance/synapse formation (DIP, DPR, SIDE, BEAT and NLG). (C) Gene ontology enrichment analysis revealed that DEGs with a p-value <0.05 are involved in biological processes such as neuron differentiation, neuron projection development, axon guidance and development. The top 15 categories are shown. (D) Gene ontology enrichment analysis revealed that DEGs with a p-value <0.05 have molecular functions such as neuron differentiation, neuron projection development, axon guidance and development. The top 15 categories are shown.

### Drgx regulates the transcription of the T4/T5 late specifier *Sox102F*

This study demonstrates the fundamental involvement of Drgx in T4/T5 neuron specification and maturation. Given the importance of Drgx in these processes, the position of Drgx in the T4/T5 transcription factor hierarchy was of interest. The observed phenotypes resemble those documented for the loss of transcription factors Ato, Dac, SoxN, and Sox102F (Contreras et al., 2018; Schilling et al., 2019). The RNA-seq dataset demonstrated a slight but non-significant reduction in expression following *Drgx* knockdown for ato, Dac, and SoxN, whereas a log2fold change of 2.04 was revealed for Sox102F. Two Drgx binding sites are present at the *Sox102F* locus, with one located directly at the TSS of the most prominently expressed spliceform in T4/T5 neurons, and the other located 8 kb downstream of the TSS (Fig. 7A). In controls, GFP expression from a MiMIC insertion at the *Sox102F* locus was visible in differentiating T4/T5 neurons (Fig. 7B, Fig. 7D). However, after *Drgx* knockdown, GFP expression was significantly (p=0.0016) reduced (Fig. 7C, Fig. 7D), thus supporting the RNA-seq results.

**Fig. 7:**
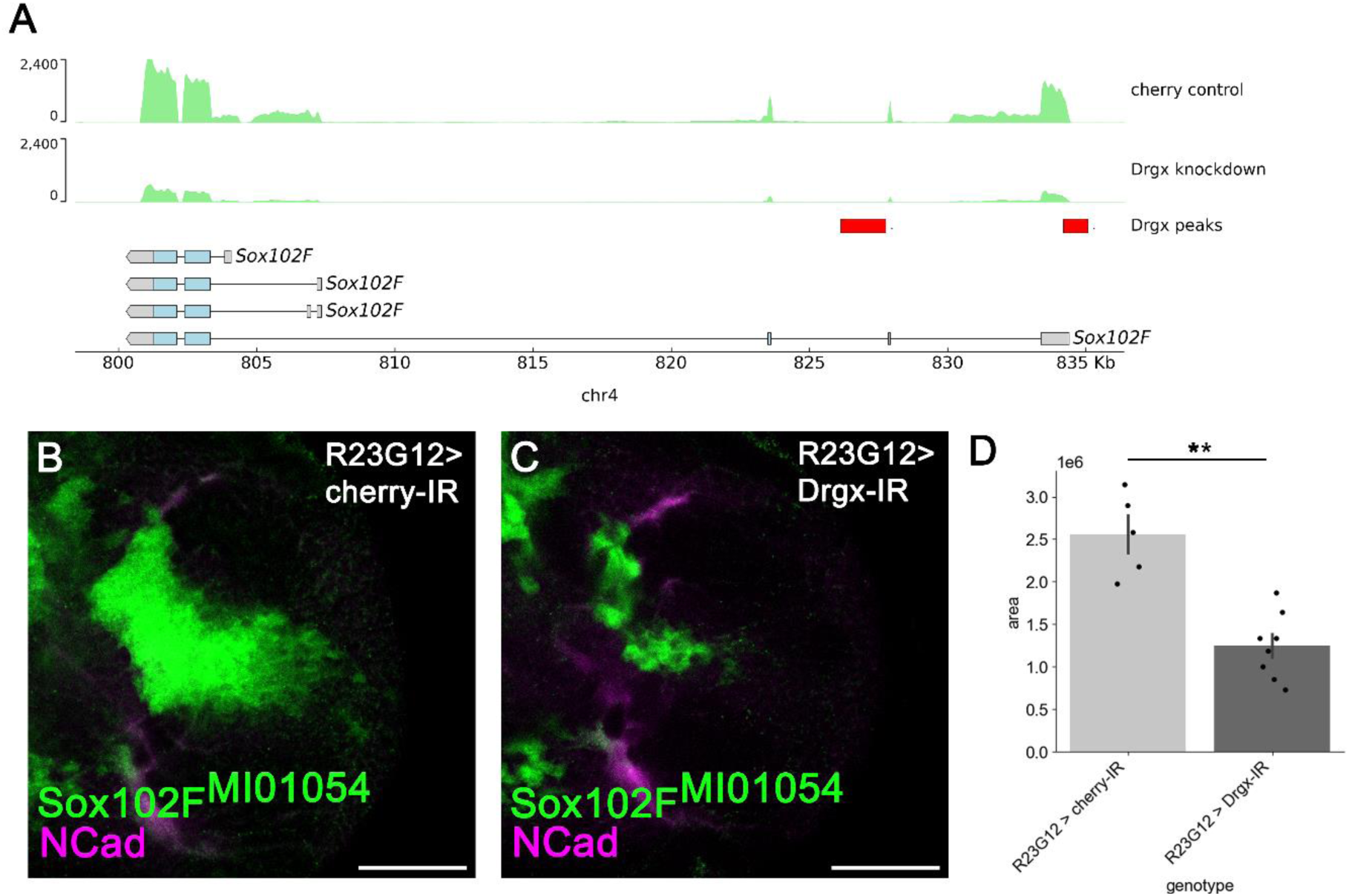
Drgx regulates the expression of *Sox102F* in the developing T4/T5 neurons. (A) In the TaDa dataset of T4/T5 neurons, two binding peaks can be observed at the *Sox102F* locus. In addition to that, the T4/T5 specific RNA-seq revealed a downregulation of Sox102f upon *Drgx* knockdown. (B) Sox102F expression is present in the developing T4/T5 neurons of the larval brain in the control (R23G12-GAL4; Sox102F^MI01054^). (C) In the larval brain after *Drgx* knockdown (UAS-Drgx-IR; R23G12-GAL4; Sox102F^MI01054^), the expression of Sox102F is significantly diminished. (D) When comparing the area of Sox102F expression in the control group (n=5) with that in the larval brain of *Drgx* knockdown (n=7), a significant (p= 0,0016) reduction is observed in the visual representation. For statistical analysis, Mann-Whitney U-Test was performed and the graph shows ± SEM. Scale bars: 50 µm.

**Fig. 8:**
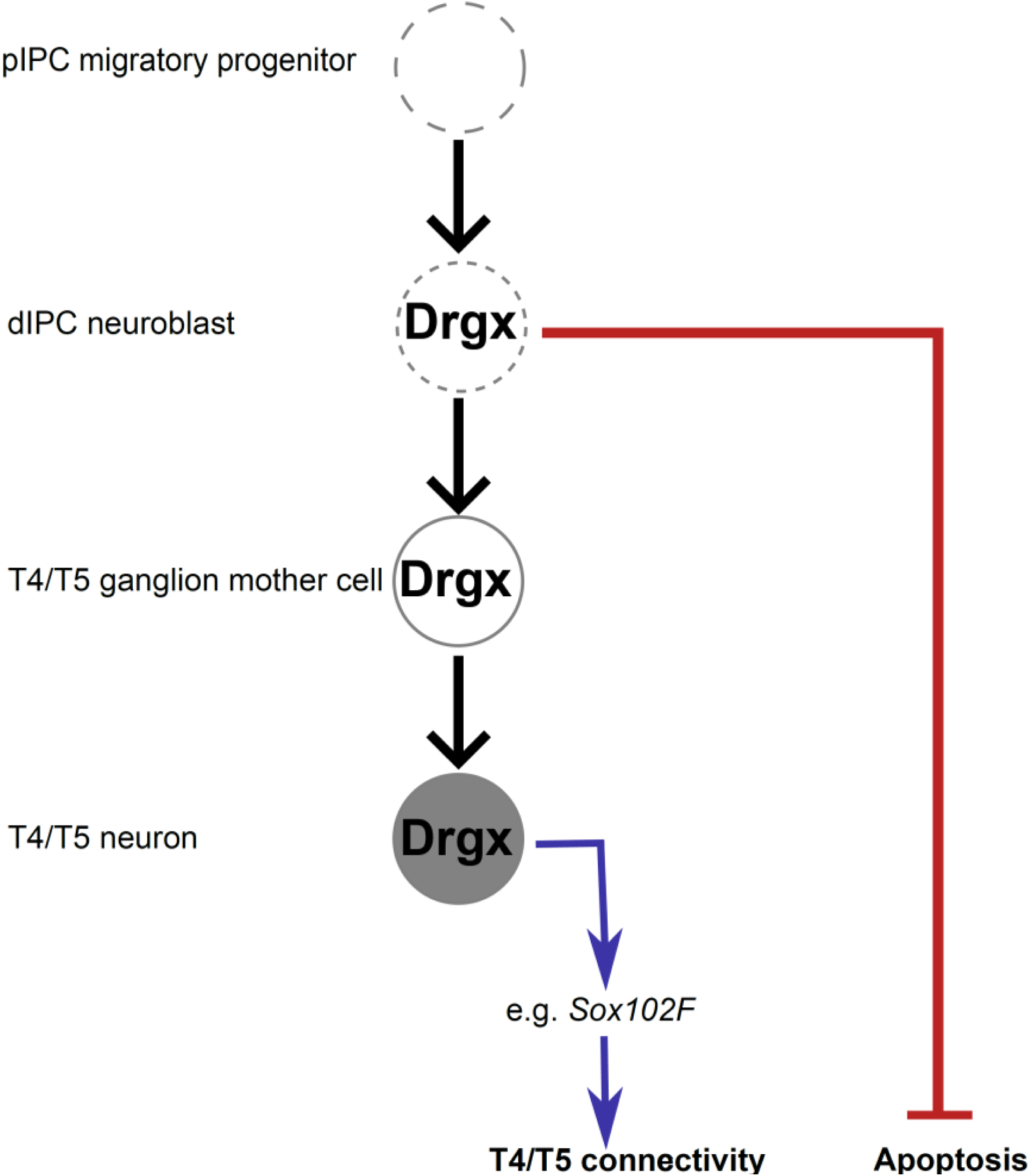
Drgx has an early and a late function during the T4/T5 neuronal development. Drgx expression can be detected in dIPC neuroblasts where its first function is to inhibit apoptosis of T4/T5 neurons by regulating specific apoptosis genes (marked in red). Later, during T4/T5 neuron development, Drgx second function is to affect the formation of the correct axon connection by influencing the expression of many axon guidance genes, e.g. Sox102F (marked in blue).

## Discussion

The typical circuitry pattern of T4/T5 neurons heavily relies on the activation of specific transcription factors. In recent years, this genetic cascade has been extensively studied and genes such as *dpp*, *dac*, *SoxN*, and *Sox102F* are found crucial for the accurate development of T4/T5 neurons (Apitz and Salecker, 2015; Apitz and Salecker, 2018; Konstantinides et al., 2022; Pinto-Teixeira et al., 2018; Schilling et al., 2019). In this study, we describe Drgx as a further essential transcription factor with a functional importance during early and late development of T4/T5 neurons. Our expression analysis revealed its presence in both larval (Fig. 1E, Fig. 1F) and adult stages (Fig. 1G) of T4/T5 neurons. We could prove that *Drgx* is expressed in dividing cells by an overlap of pHH3 staining (Fig. 2B). Furthermore, by co-staining with Dac antibody, we showed that Drgx is already expressed in T4/T5 neuroblasts of the d-IPC within the larval brain (Fig. 2A). Since Apitz and Salecker (2015, 2018) have shown that Dac is present in T4/T5 neuroblasts of the d-IPC, where it, together with Ato, is essential for the proper formation of mature T4/T5 neurons, it can be used as a T4/T5 neuroblast marker. We concluded that Drgx plays an important role in the initial development of T4/T5 neurons.

In the motion-sensing visual pathway, T4/T5 neurons constitute the first stage of computation, and the optomotor response is highly dependent on their proper functioning. This was shown by blocking these neurons with a temperature-sensitive shibire mutation, resulting in a complete loss of the optomotor response in flies (Bahl et al., 2013; Schnell et al., 2012). Our optomotor experiments revealed comparable results when inducing pan-neuronal and T4/T5 neuron-specific knockdowns of *Drgx* showing that the knockdown of *Drgx* significantly affects the performance of T4/T5 neurons in *Drosophila* (Fig. 3). By conducting the Buridan’s paradigm test, we were able to eliminate the prospect of *Drgx* knockdown flies being completely blind (Fig. S2). Consequently, it was deduced that *Drgx* expression in neurons mostly affects the motion detection ability of the fly’s visual system.

The mistargeting of T4/T5 dendrites and axons observed upon *Drgx* knockdown in the pupal (Fig. S3) and adult brain (Fig. 4) in this study is akin to the phenotypes detected in *dac* and *ato* mutants, whereby the T4/T5 neurons also mistarget their intended stopping layers in the lobula and medulla (Apitz and Salecker, 2018). Furthermore, Apitz and Salecker (2018) also observed a similar reduction in lobula plate size in double *ato* and *dac* knockdown flies as we observed in *Drgx* knockdown flies. Finally, Apitz and Salecker (2018) showed that T4/T5 neuron identity is lost in the simultaneous knockdown of *ato* and *dac* in d-IPC neuroblasts. All these results, together with the early expression of Drgx in d-IPC neuroblasts simultaneously with Dac (Fig. 2A), suggest that Drgx plays a significant role in regulating T4/T5 neuron identity and circuit formation of these neurons. This is consistent with the observation of a similar effect in mice with the Drgx orthologue, DRG11. Several studies have shown that dorsal root ganglia neurons in mutant mice are unable to innervate spinal gray matter, resulting in reduced sensitivity to noxious stimuli (Chen et al., 2001; Rebelo et al., 2007; Rebelo et al., 2010). Although the phenotypes are very similar, in this study we could not demonstrate a direct regulation between Dac, Ato and Drgx. At present, our results suggest that these are rather parallel but independent pathways.

Another specifier of T4/T5 identity that was recently described in two independent publications is Sox102F (Contreras et al., 2018; Schilling et al., 2019). Both Contreras et al. (2018) and Schilling at al. (2019) showed that *Sox102F* mutant and knockdown flies have defects in terminal differentiation and proper targeting of T4/T5 dendrites. The phenotypes observed in these publications are the same as those we observed upon *Drgx* knockdown, such as loss of optomotor function due to overgrowth of T4/T5 dendrites in the medulla and lobula together with a reduced size ratio between the lobula plate and medulla. In contrast to Drgx expression, Sox102F expression was not found in T4/T5 neuroblasts making Sox102F a late T4/T5 neuron specifier (Contreras et al., 2018; Schilling et al., 2019). This, together with the Drgx binding site in the *Sox102F* gene and the downregulation of *Sox102F* by *Drgx* knockdown, suggests that *Drgx* is positioned above *Sox102F* in the developmental gene cascade and acts as an early T4/T5 neuronal specifier upstream of Sox102F.

The early positioning of Drgx in the gene cascade of T4/T5 neuron development is also supported by the increased apoptosis observed in the developing larval optic lobe (Fig. 4I, Fig. 4K), which leads to a reduced number of T4/T5 neurons. In our experiments, the increase in apoptosis was not only demonstrated histologically, but our RNA-seq also revealed the upregulation of known apoptosis markers, including *reaper* (*rpr*) (White et al., 1994), *grim* (Chen et al., 1996) and *sickle* (*skl*) (Christich et al., 2002). In DRG11 null mutant mice, a similar increase in apoptosis and consequent reduction in the number of neurons was observed in dorsal root ganglia neurons. It has been hypothesized that these neurons die because they are unable to make the proper connections (Rebelo et al., 2006). This need for neurons to connect with other neurons to survive was described decades ago (Levi-Montalcini, 1987; Oppenheimer, 1991).

Although the mistargeting effect seen in *Sox102F* mutants and knockdowns is like that observed for *Drgx*, no increase in apoptosis was observed (Contreras et al., 2018). While this seems contradictory at first, it should be noted that our observation of increased apoptosis only occurred when using the early T4/T5 driver lines R9B10-GAL4 or R23G12-GAL4, but not with the late driver R39H12-GAL4, which only shows the mistargeting effect upon *Drgx* knockdown. This suggests that Drgx is also essential for the early development of T4/T5 neurons before it activates the expression of *Sox102F*. Thereby supporting the idea that Drgx has a dual role in the development of T4/T5 neurons. First, it is essential for the survival of the neuron, and later it is needed to form the correct neuronal connections.

The formation of neuronal connections depends on the expression of a large number of different genes. In both the TaDa analysis and the RNA-seq, in the GO enrichment analyses among the top 15 categories, terms such as cell projection morphogenesis, axon guidance, axon development and synapse organization can be found (Fig. 5D, Fig. 5E, Fig. 6C, Fig. 6D). This again highlights the significant role Drgx plays in the development of the correct T4/T5 neuron connectivity. Additionally, especially the RNA-seq showed that the *Drgx* knockdown did not only lead to the up- or downregulation of single axon guidance genes but rather to the different regulation of whole axon pathfinding families like the DIP/dpr, neurolin (nlg) and the sidestep (side)/ beaten path (beat) family (Fig. 6B). Therefore, a completely different expression profile is created upon *Drgx* knockdown, causing an alteration in the circuitry development, which ultimately leads to an identity loss of the T4/T5 neurons.

In conclusion, Drgx acts as a key regulator of T4/T5 neuron identity and is required together with Dac, Ato and Sox102F for the specification of T4/T5 neurons. During early T4/T5 development, Drgx is essential for the survival of developing T4/T5 neurons and later controls axon guidance and the formation of the correct dendritic arborization of T4/T5 neurons in the medulla and lobula. This is achieved through the transcriptional regulation of several genes involved in pathfinding belonging to the DIP/dpr, Nlg and Side/Beat families and, most importantly, *Sox102F*.

## Material and Methods

### Fly strains

All flies were maintained on a 12-hour day-night cycle at 25 °C and 65% humidity unless otherwise noted. As a food source, standard cornmeal agar medium was provided. The following stocks were obtained from the Bloomington Drosophila Stock Center (BL): y[1]w[*]; P{w[+mC]=Act5C-GAL4}17bFO1/TM6B,Tb[1] (BL#3954), P{w[+mW.hs]=GawB}elav[C155] (BL#458), y[1]w[*]; Mi{y[+mDint2]=MIC}Drgx[MI06689] (BL#42422), y[1]w[*]; Mi{y[+mDint2]=MIC}Drgx[MI11472]/SM6a (BL#56323), w[1118]; P{y[+t7.7]w[+mC]=GMR40E11-GAL4}attP2 (BL#48140), w[1118]; P{w[+m*]=GAL4}repo/TM3,Sb[1] (BL#7415), w[*]; P{w[+mC]=tubP-GAL80[ts]}20; TM2/TM6B,Tb[1] (BL#7019) w[*]; P{y[+t7.7] w[+mC]=10XUAS-IVS-mCD8::GFP}attP40 (BL#32186), w[*]; P{y[+t7.7]w[+mC]=15XUAS-IVS-mCD8::GFP}attP2 (BL#32193), y[1]sc[*]v[1]sev[21]; P{y[+t7.7]v[+t1.8]=UAS-mCherry.VALIUM10}attP2 (BL#35787), y[1]sc[*]v[1]sev[21]; P{y[+t7.7] v[+t1.8]=TRiP.HMC04912}attP40 (BL# 57723), WTB (BL#8522), w[*]; P{w[+mC]=13xLexAop2-CD4-tdTom}4/CyO, P{Wee-P.ph0}Bacc[Wee-P20] (BL#77138). The rest of the stocks were either created during the course of this study or gifted by fellow researchers: y[1]w[*]; Mi{mCherry-SV40.2}Drgx[MI11472-mCherry-SV40.2]/CyO (created via RCME), y[1]w[*]; Mi{GAL4-Hsp70pA.2} Drgx[MI11472-GAL4-Hsp70pA]/CyO (created via RCME), y[1]w[*]; Mi{GAL4-Hsp70pA.2} Drgx[MI11472-GAL4-Hsp70pA]/CyO (created via RCME), w[1118]; P{y[+t7.7]w[+mC]=GMR23G12-GAL4}attP2 (gifted by Tabea Schilling), w[1118]; P{y[+t7.7]w[+mC]=GMR39H12-GAL4}attP2 (gifted by Tabea Schilling), w[1118]; P{y[+t7.7]w[+mC]=GMR9B10-GAL4}attP2 (gifted by Tabea Schilling), w[*]; P{w[+mC]=repoGal80}attP86Fb (gifted by Christian Klämbt), w[*]; UAS-LT3-NDam (gifted by Andrea H. Brand), w[*]; UAS-LT3-NDam-Drgx (created in our lab), CG14340-LexA (created in our lab).

### Embryonic microinjections

For the injection solution, 150ng of the vector (in Ampuwa H_2_O) and 10x Injection Buffer (25mM NaCl, 5mM KCl, 0.1mM EDTA in H_2_O) were diluted in Ampuwa H_2_O, centrifuged at maximum speed for 4min and transferred to a microinjection glass capillary needle. Prior to the day of injection, the flies were placed on apple agar plates (33.3g Sucrose, 330mL Apple juice, 26.7g Bacto-Agar, 1000mL H_2_O, 20ml 10% Nipagin) with a yeast suspension on top. While injecting, they were placed on new plates every 30min to ensure the proper age of the embryos. The embryo collection and injection took place at 18°C.The embryos were washed from the agar plates and their chorion was removed by rolling them on double-sided tape before placing them on coverslips, drying them with silica gel for 10min, and covering them with oil. The prepared mixture was injected into the pole cells of the embryo using compressed N_2_. Embryos were then placed on small apple agar plates to complete their development at 18°C during the next two days. The hatched larvae were collected and placed in food vials at 25°C until the adult flies’ eclosed. These flies were then crossed with the correct balancer strain, and positive transformants were selected in the next generation of flies. The following vectors of the Drosophila Genomics Resource Center were used for injection: pBS-KS-attB1-2-GT-SA-GAL4-Hsp70pA (1325), pBS-KS-attB1-2-GT-SA-mCherry-SV40 (1324) and pBS-KS-attB1-2-PT-SA-SD-2-EGFP-FIAsH-StrepII-TEV-3x-Flag (1314). Additionally, the cloned vector pNDam-DRGX3 was also injected.

### Optomotor assay

The optomotor response of walking flies was tested by placing the flies on a movable platform in the middle of an illuminated arena. During the experiment, black stripes moved along the illuminated walls of the arena. The movement of the platform was transferred to a computer via a photoelectric sensor and was analyzed using a python-based script (von der Linde, 2018). Before the experiment, a small piece of fishing line was glued to each flýs thorax during cold anesthesia, thereby ensuring that the fly is able to move its head and abdomen. The flies were tested for 5min whereby the rotation of the black stripes changed every 30 seconds. The results were analyzed with the R based script DTSevaluation available at https://github.com/brembslab/DTSevaluations.

### Buridańs paradigm

Spatial landmark orientation was tested in three-to-four-day old male flies. One day prior to testing, their wings were clipped under CO_2_ anesthesia and until testing they were placed in individual food vials. The walking behavior was tested in a uniformly illuminated cylinder with a platform in the middle, which was surrounded by water. On opposing sides of the cylinder, black stripes were the only landmarks for the flies. Each clipped fly was placed on top of the platform and walked freely for 15min. The walking path of the fly was recorded with a camera, which was controlled by the BuriTrack software (http://buridan.sourceforge.net). The results were analyzed using CeTrAn (https://github.com/jcolomb/CeTrAn).

### Histology and imaging

Wandering L3 larval, pupal and adult brains were dissected in ice cold PBS and fixed in 4% paraformaldehyde in PBS for 20min at room temperature. Before adding the primary antibodies, the brains were washed three times with PBS. The following day after the primary antibody staining, the brains were washed again with PBS and the secondary antibodies were added. All antibodies were diluted in 0.1% PBST (PBS with 0.1% Triton X-100) with 5% normal goat serum (NGS) and stainings were carried out overnight at 4°C. After staining, the brains were washed with PBS and mounted in Vectashield (VECTOR Laboratories). The following antibodies were used: rat anti-DN Cad (1:20, DN-Ex #8, Developmental Studies Hybridoma Bank), mouse anti-Dac (1:20, mAbdac 1-1, Developmental Studies Hybridoma Bank), chicken anti-GFP (1:1000, A10262, Invitrogen/ Thermo Fisher), rabbit anti-pHH3 (1:200, Ser10, Sigma-Aldrich/Merck), mouse anti-LacZ (1:20, JIE7, Developmental Studies Hybridoma Bank), rat anti-RFP (1:500, 5F8, Chromotec), mouse anti-NC82 (1:20, stock collection), mouse anti-Acj6 (1:20, Developmental Studies Hybridoma Bank), rat anti-Elav (1:50, Developmental Studies Hybridoma Bank), mouse anti-Repo (1:20, 8D12, Developmental Studies Hybridoma Bank), goat anti-chicken-Alexa Fluor488 (1:200, Invitrogen/Thermo Fisher), goat anti-mouse-Alexa Fluor555 (1:200, Invitrogen/Thermo Fisher) and goat anti-rat-Alexa Fluor633 (1:200, Invitrogen/ Thermo Fisher), anti-rabbit-Alexa Fluor555 (1:200, Invitrogen/ Thermo Fisher). The mounted brains were imaged using the Leica TCS SP8 with the appropriate laser lines for the fluorophore-coupled secondary antibody. Leica LAS X software was used to acquire the images and the images were later processed using Fiji ImageJ software.

### Paraffin sections

Flies were anesthetized with CO_2_ and placed in a collar. The desired genotypes were positioned in an alternating manner with *sine occulis* mutant flies for orientation. The collars were then placed in Carnoy solution for 3.5 hours at room temperature. Afterward, the collars were washed three times for 30 minutes in ethanol and left overnight in methyl-benzoate. The following day, a mixture of 15 ml of paraffin and 15 ml of methyl-benzoate was prepared and incubated at 60°C for 15-20 minutes. The collars were then incubated in the mixture for 1 hour at 60°C. Subsequently, the collars were placed in paraffin for 30 minutes, with the paraffin being changed six times. After washing, the collars were placed in a plastic frame, which was then filled with paraffin and allowed to cool on a cooling platform. The cooled paraffin block was then placed on a metal block and removed from the frame. The heads are retained in the paraffin block after removing the collar. The block is then cut into shape and paraffin sections are produced on a microtome. These sections are placed on glass slides coated with eggwhite-glycerine and water. The slides are heated for two minutes to allow the sections to stretch slightly. After removing excess water, the slides are labeled and left to dry overnight. To eliminate paraffin from the sections, the slides are immersed in Xylene twice for 20 minutes at room temperature. Finally, the slides are left to dry overnight. Subsequently, the slides are incubated in Xylene for 1 hour at 60°C. After incubation, excess Xylene is removed with a tissue and the slides are sealed with coverslips using DePeX.

### Lobula plate measurements

Paraffin sections of fly heads with the desired phenotype were imaged using a fluorescent microscope. The area of the lobula plate was then measured using the Measure tool in Fiji ImageJ software and divided by the area of the medulla, which served as a reference.

### Sox102F fluorescent measurements

The total fluorescence of the Sox102F signal was calculated using open-source software Fiji ImageJ, following the protocol found at https://theolb.readthedocs.io/en/latest/imaging/measuring-cell-fluorescence-usingimagej.html. To ensure comparability, the same image plane was used for each larval brain. Finally, the measurements for the control and the knockdown were compared.

### TaDa sample acquisition

TaDa crosses were kept at 18°C until L3 larvae stage. After a 24 hours induction at 30°C, 30 wandering L3 larvae were collected per genotype, and their brains were dissected in cold PBS for one hour before flash freezing them in liquid N_2_. For the gDNA extraction and sample preparation the protocol of Marshall et al. (2016) was used. To reduce DNA fragment size, the sonication step was performed with 6 cycles of 30s on and 90s off. The libraries were prepared using the NEBNext Ultra DNA Library Prep Kit for Illumina together with the NEBNext Multiplex Oligos for Illumina according to the manufacturers protocol. The finished libraries were in the end analyzed with an Agilent 4200 TapeStation to ensure a medium fragment length between 400 to 450bp.

### TaDa Sequencing and Bioinformatic analysis

The libraries were sequenced at the NGS (next generation sequencing) Core-Facility of the Leibnitz-Institut für Immuntherapie (LIT) on a NextSeq 550 (Illumina) sequencer. Single end sequencing was performed with 20 million reads per library and 10% PhiX spike-in. Before analyzing the obtained data, uncut DamID adaptors and reads containing an initial or internal GATC site were removed with fastq-filter (github.com/owenjm/damid_misc) followed by seqkit (Shen et al., 2016). Afterwards the data was analyzed using the damidseq_pipeline from Marshall and Brand (2015). This involved aligning the sequenced DNA fragments to the *Drosophila* genome BDGP6.32, read extension, binned counts, normalisation, pseudocount addition and final ratio file generation. For peak calling, bam files with extended reads for Dam only controls or Drgx-Dam fusion samples generated by the pipeline were merged with samtools (Li et al., 2009). Peaks were called with find_peaks (https://gist.github.com/efekarakus/cc7303456841523f37dd) and min_quant set to 0.9. Datasets were visualized using the igv-browser or pygenometracks (https://academic.oup.com/bioinformatics/article/37/3/422/5879987). Averaged Drgx-Dam/Dam signal across pseudogenic region and heatmap of DamID was visualized with SeqPlots. Drgx peak count frequency was visualized with ClusterProfiler. Distribution of Drgx peaks was analyzed with ChIPpeakAnno. Gene ontology analysis was performed using the clusterProfiler R package. The Benjamini-Hochberg multiple testing correction method was used and adjusted *P*-value cutoff was set to 0.01.

### Single-cell suspension and cell sorting

To establish a protocol for creating the single-cell suspension Li et al. (2017) and Hörmann et al. (2020) were used as a template. Wandering L3 larval brains (cherry-IR n=107, Drgx-IR n=114) with fluorescently labeled T4/T5 neurons were dissected in DSM (Drosophila Schneiders medium, PAN Biotech) for one hour. During this time, the finished brains were placed on ice in DSM. Prior to dissection, a fresh 100 units/mL papain (Worthington) solution was prepared. After dissection, the brains were washed three times with DSM. Between each washing step, the brains were centrifuged at 2600 rpm for 1min at 4°C. The dissociation solution was then added to the brain (157.9µL DSM, 50µL papain solution (100units/mL) and 2.1µL Liberase (2.5mg/mL, Roche)) and dissociation took place for 30min at 30°C. Digestion was stopped by adding 300µL of ice-cold DSM. The brains were again washed three times with DSM at 2600rpm for 1min at 4°C. Finally, 600µL of sterile PBS containing 1% BSA (Bovine serum albumin, Pan Biotech) was added to the brains and the brains were mechanically disrupted by being pipetted up and down with a 200µL pipette. To remove any cell clumps, the solution was strained through a 40-µm Flowmi filter into a FACS tube and stored on ice until sorting. Sorting took place on a BD FACSAria IIu of the LIT Central FACS Facility (CFF). Propidium iodide (PI, Merck) was used to identify dead cells and was added to the cells directly prior to sorting at a final concentration of 1ng/mL. Cells were sorted directly into the β-mercaptoethanol supplemented RTL Plus buffer of the RNeasy Plus Micro Kit (Qiagen).

### RNA extraction and Library preparation

The RNA was extracted from the sorted cells using the RNeasy Plus Micro kit according to the manufacturer’s protocol. The cells were lysed using the QIAshredder and gDNA was removed with gDNA eliminator spin columns provided with the RNeasy Plus Micro kit. The RNA libraries were prepared with the SMART-Seq v4 PLUS Kit from TaKaRa following the TaKaRa protocol. For cDNA synthesis, 2µL of 3’SMART-Seq CDS primer was used for first-stand synthesis and cDNA was amplified with 10 PCR cycles. The finished cDNA sample was measured at the 4200 TapeStation using the HS D5000 ScreenTape. For the final library preparation, the maximum amount of cDNA was used, and the last amplification PCR was performed with 15 cycles. At last, the concentration of the finished samples was measured with the Qubit dsDNA-HS assay kit on the Qubit 3 Fluorometer and the fragment length distributions were measured with the D5000 ScreenTape on the 4200 TapeStation.

### RNA library sequencing and bioinformatic analysis

The sequencing was again performed at the LIT NGS Core-Facility on a NextSeq 550 (Illumina) sequencer performing single end sequencing. Thereby, 30 million reads per library were generated. The bioinformatic analysis was performed using the following programs: STAR 2.7.10a (Dobin et al., 2013) with --quantMode GeneCounts was used to align reads to the Drosophila transcriptome (BDGP 6.32) and to generate count matrices from each library. Subsequently, sam files were converted to bam files, sorted and indexed using samtools (Li et al., 2009). Bam files were converted to bigwig files using bamCoverage from the deepTools package (https://github.com/deeptools/deepTools) for visualization in the igv-browser or in pygenometracks. Gene count matrices from different libraries were combined into a single matrix in R, and DEseq2 (Love et al., 2014) was used to perform differential gene expression analyses and examine data quality. Genes without an unique Flybase gene ID were excluded from these analyses. R package EnhancedVolcano was used to generate volcano plot and pheatmap to generate heatmap. Gene ontology analysis was performed with DEGs with an adjusted p-value <0.05 using the clusterProfiler R package. The Benjamini-Hochberg multiple testing correction method was used and adjusted p-value cutoff was set to 0.01.

### Statistical analyses

All statistical analysis were performed using the python library SciPy and bar plots display mean ± SEM. A two-sided Mann-Whitney U-Test was conducted to determine statistical significance. The p-values are displayed as *p-value ≤0.05, **p-value ≤0.01, ***p-value ≤0.001 in all figures.

## Data Availability

The TaDa and RNA-seq datasets generated and analyzed as part of this study are available under GSE253077 and GSE253078. The data on which the diagrams are based can be found in the supplementary data.

## Acknowledgements

We would like to thank Tabea Schilling, Christian Klämbt and Andrea H. Brand for the provided flies. For the cell sorting, we want to thank Rüdiger Eder and Petra Hoffmann from the LIT Central FACS Facility (CFF). We would also like to thank Michael Rehli, Claudia Gebhard, Nicholas Strieder and Hanna Stanewsky from LIT NGS Core-Facility for the library sequencing. Further thanks also go to Björn Brembs for access to his optomotor and Buridańs paradigm equipment. We would also like to thank Renate Reng for her help with the microinjections of the *Drosophila* embryos. In addition, a big thanks to Svenja Oestreich for her help dissecting the larval brains for RNA sequencing. Finally, we would also like to thank the Bloomington Drosophila Stock Center, the Developmental Studies Hybridoma Bank and the Drosophila Genomics Resource Center for providing us with various flies, antibodies and vectors.

## Supplemental Figures

**Supplemental Figure 1:**
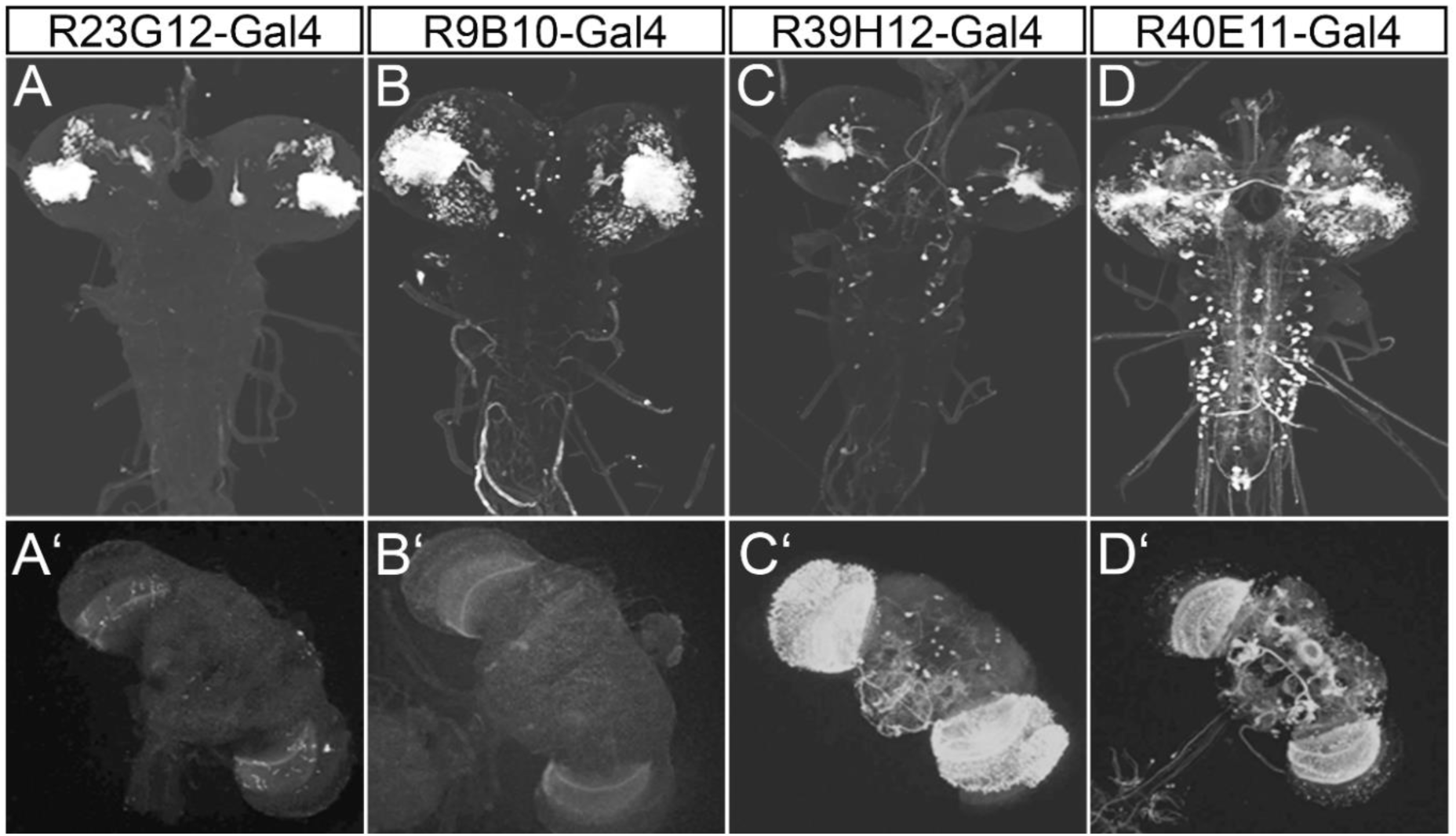
Expression pattern of the T4/T5 FlyLight enhancer GAL4 lines used in this work. The GAL4 enhancer lines that functioned as T4/T5 neuron-specific driver lines varied in both expression pattern and expression strength. (A, A’) Among them, R23G12-GAL4 was the most frequently useded line due to its (A) strong expression in larvae and (A’) moderate expression in adults. (B, B’) The expression pattern of R9B10-GAL4 is comparable to the first line, (B’) yet it is significantly weaker in adult brains. (C, C’) R39H12-GAL4 expression initiates at a later stage and (B) is limited to a small number of T4/T5 neurons in L3 larva. (B’) However, in the adult brain, the line exhibits strong expression in T4/T5 neurons, along with the expression in other cells in the optic lobe besides the T4/T5 neurons. (D, D’) The R40E11-GAL4 line has the broadest expression pattern, encompassing neurons in both the larval ventral nerve cord and adult central brain. Images utilized are from the FlyLight website of the Janelia Research Campus (Jenett et al., 2012). To reflect their expression pattern, all GAL4 lines were crossed with UAS-CD8-GFP.

**Supplemental Figure 2:**
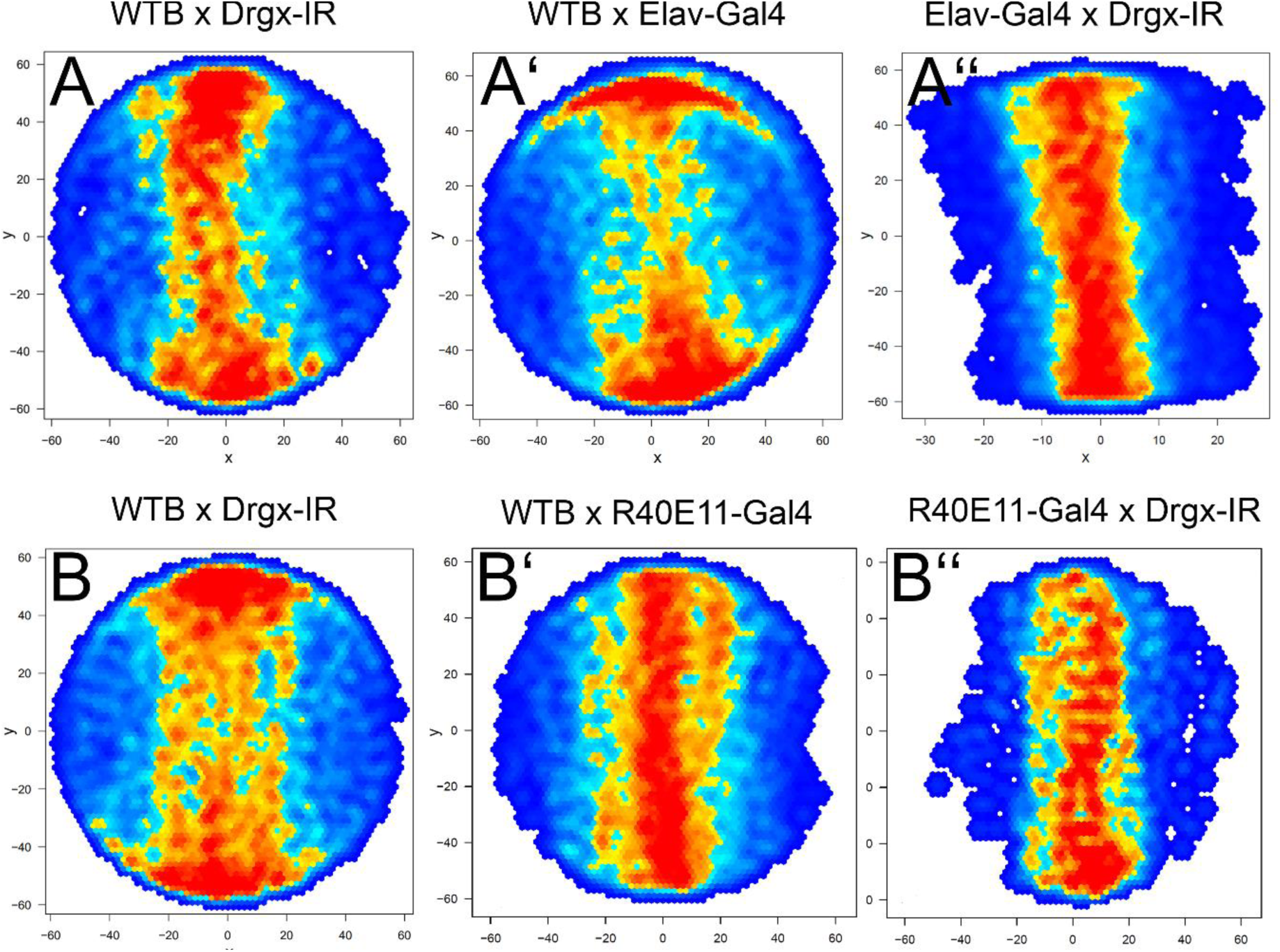
*Drgx* knockdown in T4/T5 neurons does not lead to blindness. The visual ability of *Drgx* knockdown flies was assessed using the Buridańs paradigm. (A) In the transition plot the UAS-Drgx-IR knockdown control (n=30) as well as the (A’) Elav-GAL4 control (n=33) and (A’’) the Elav-GAL4; UAS-Drgx-IR knockdown flies (n=25) show a normal walking behavior indicating that these flies are not blind. (B, B’, B’’) The same result was also observed for the T4/T5 specific *Drgx* knockdown using the R40E11-GAL4 driver line (WTBxDrgx-IR n=10, WTBxR40E11-GAL4 n=12, R40E11-GAL4xDrgx-IR n=11). The transition plots shows the relative frequency of fly passage at any position in the arena. Red indicates a high frequency whereas blue indicates a low frequency of fly passage.

**Supplemental Figure 3.**
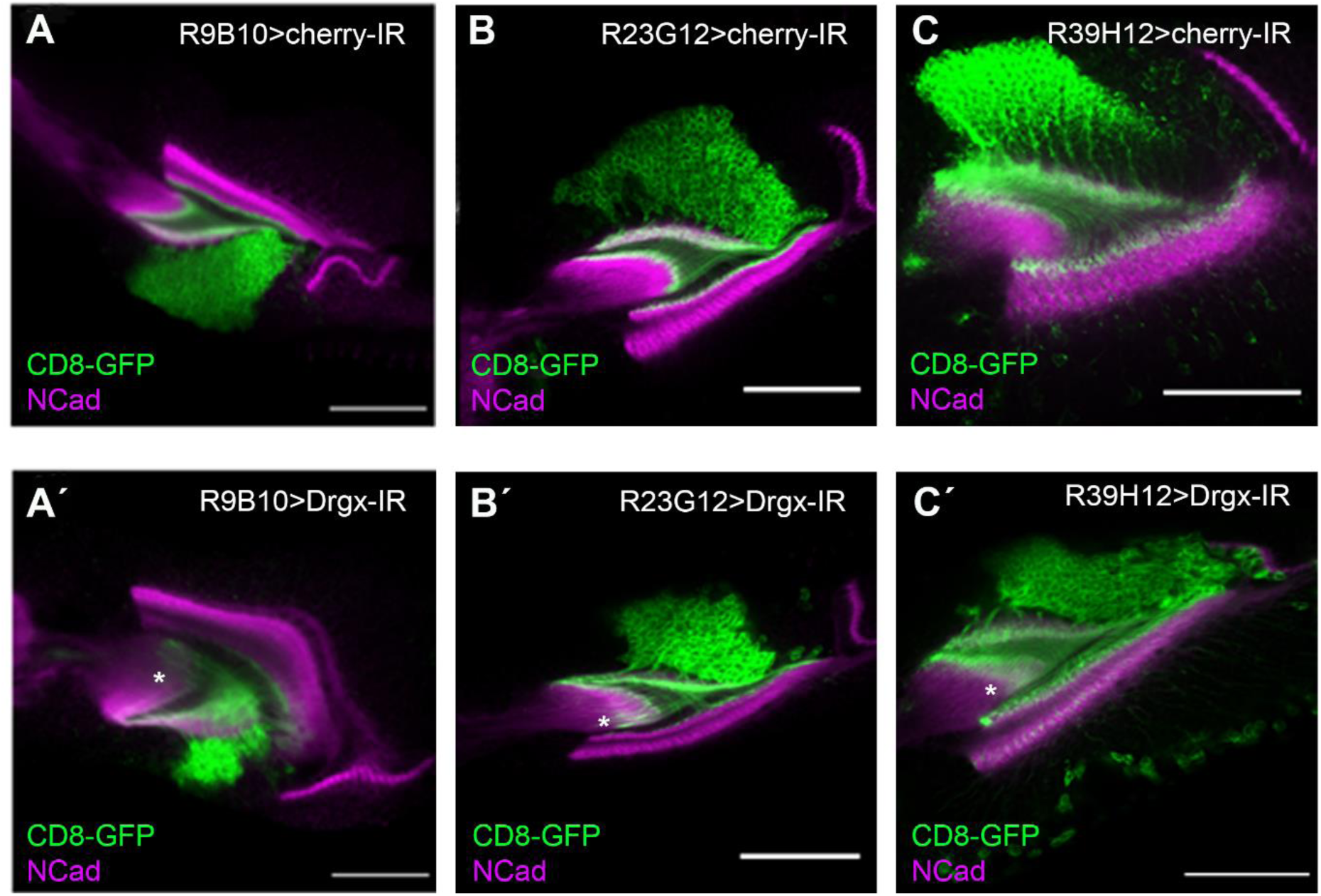
Pupal Drgx knockdown brains show T4/T5 mistargeting at 24 hours APF. (A-C) In 24 hours APF pupae, the optic lobe neuropil, labeled with anti-NCad, already exhibits a similar organization to that of the adult optic lobe, as demonstrated in the control images of all three driver lines. The driver lines used were (A) R9B10-GAL4, (B) R23G12-GAL4, and (C) R39H12-GAL4, and controls were generated by crossing these lines with UAS-CD8-GFP; UAS-Cherry-IR. (À-C’) In the *Drgx* knockdown (UAS-Drgx-IR; UAS-CD8-GFP) using again (A’) R9B10-GAL4, (B’) R23G12-GAL4 and (C’) R39H12-GAL4 the dendrites extend too far into the lobula and medulla (asterisk) resembling the phenotype observed in adult knockdown brains. Scale bars: 50 µm.

**Supplemental Figure 4.**
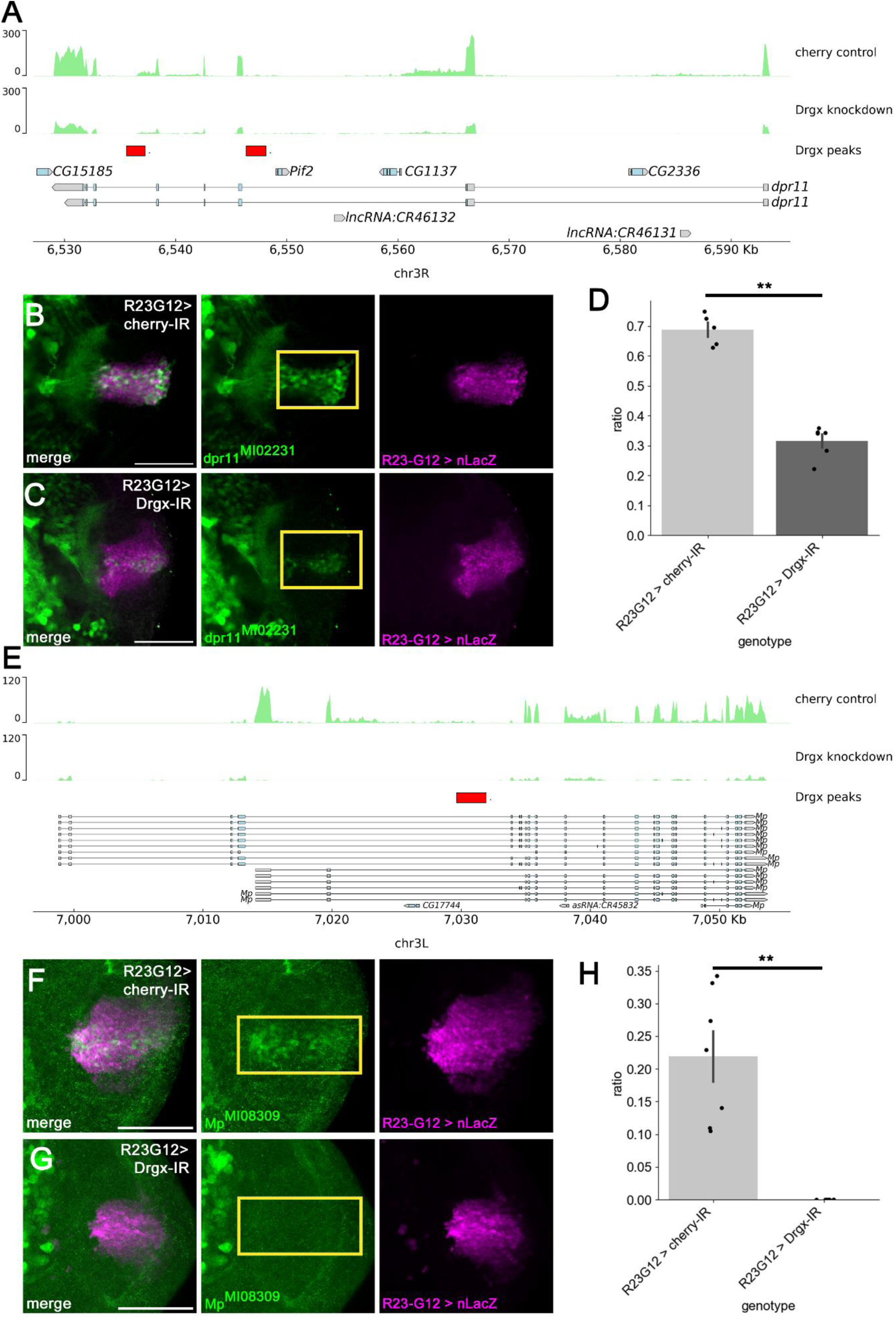
T4/T5 specific knockdown of *Drgx* reduces the expression of Dpr11 and Mp. (A) The TaDa data visualization using the IGV browser revealed two peaks at the end of the *dpr11* gene sequence and the RNA-seq data shows a reduction of *dpr11* expression upon *Drgx* knockdown. (B) MiMIC dpr11 (dpr11^Mi02231^) exhibited significant expression in T4/T5 neurons (yellow box) in wild-type flies (UAS-nLacZ; R23G12-GAL4/dpr11^Mi02231^). (C) This expression decreased (yellow box) following *Drgx* knockdown (UAS-nLacZ/UAS-Drgx-IR; R23G12-GAL4/ dpr11^Mi02231^). (D) The Dpr11 fluorescence signal was validated, revealing a ratio of 0.69 in the control group (n= 5). The knockdown group (n=6) exhibited a significantly lower ratio of 0.32 (p-value=0.0043). (E) The expression of *Mp* shows a significant reduction in the RNA-seq data upon knockdown of *Drgx*. Additionally, one peak was identified in the T4/T5 neuron specific TaDa data of the *Mp* gene sequence. (F) Fluorescent signal was only detectable (yellow box) in the brain of control larvae (UAS-nLacZ; R23G12-GAL4/Mp^MI08309^). (G) In the brain with *Drgx* knockdown (UAS-nLacZ/UAS-Drgx-IR; R23G12-GAL4/ Mp^MI08309^), no expression was observed (yellow box). (H) In the area analysis, the control exhibited a ratio of 0.22 (n=7), which was significantly higher (p-value=0.0011) than the ratio of 0 observed in the *Drgx* knockdown (n=7). All graphs display ± SEM, and statistical analysis was performed using the Mann-Whitney U test. Scale bars: 50 µm.

**Supplemental Figure 5.**
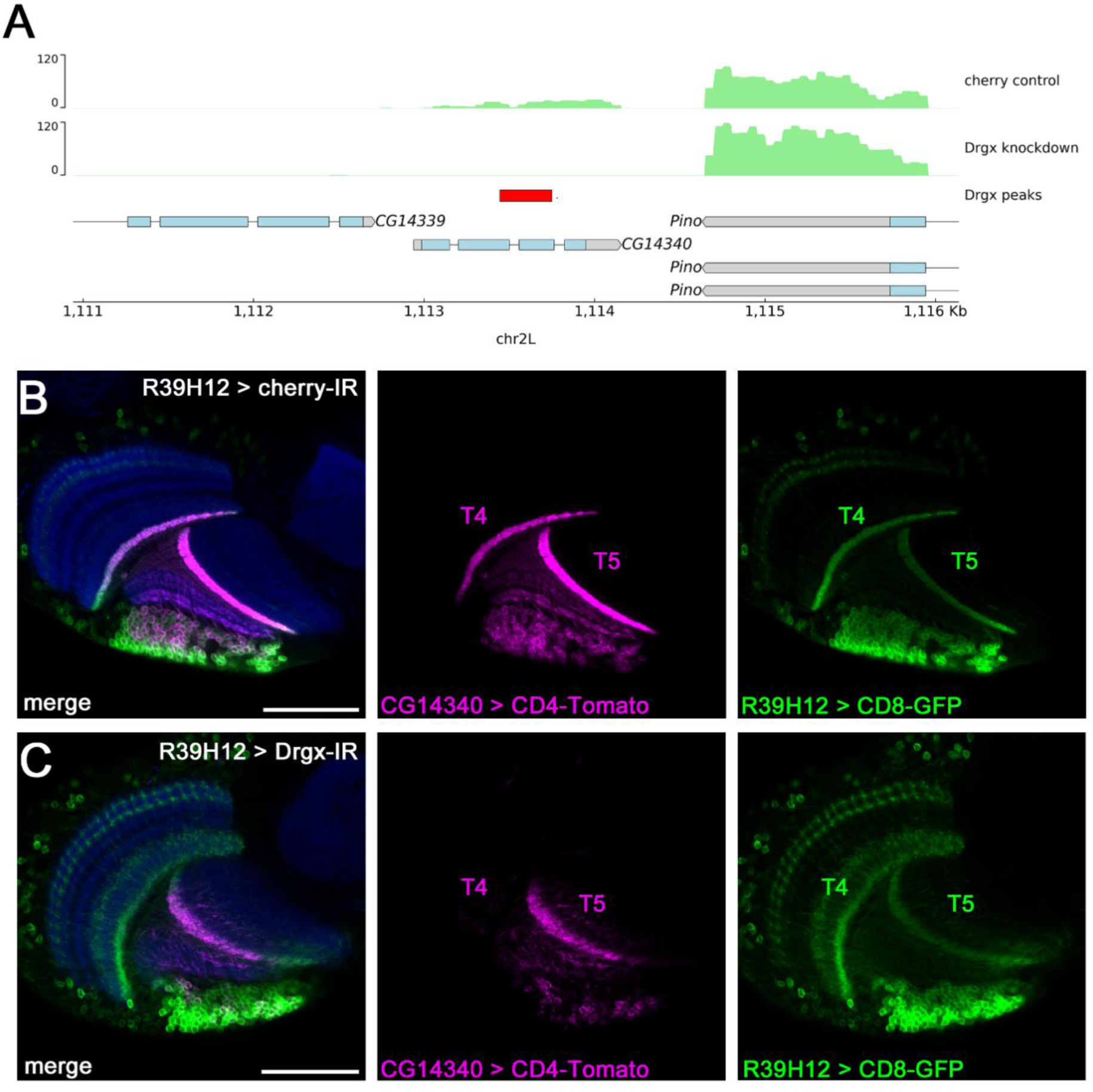
Drgx is essential for the T4 expression of CG14340. (A) The RNA-seq data set showed a complete lack of expression of CG14340 in the *Drgx* knockdown compared to the control. In the T4/T5 specific TaDa data set, a Drgx binding site was found in the CG14340 gene. (B) CG14340 is expressed in all T4 and T5 neurons of control flies (CG14340-LexA, LexAop-CD8-RFP; R39H12-GAL4, UAS-CD8-GFP). (C) Knockdown of *Drgx* in T4/T5 neurons completely abolishes CG14340 expression in T4 neurons and significantly reduces it in T5 neurons. Additionally, the dendritic overgrowth phenotype specific to the *Drgx* knockdown can also be observed. Scale bars: 50 µm.

## Supplemental Method

### Quantifying relative expression of Dpr11 and Mp based on fluorescence

To quantify changes in gene expression after Drgx knockdown using fluorescence signal, we calculated the ratio between the area of the target gene expression and the area of the cells in which it is expressed. We used the measure plugin in the open-source software Fiji ImageJ for this purpose. To measure the expressing cells’ area, we traced it using the freehand selection tool and calculated the enclosed area size with the measure plugin. We repeated the same process for the target gene expressing area and then divided the area size of the target gene expressing area by the area size of the expressing cells. We performed this procedure on all control and knockdown larval brain pictures. Ensure that measurements are taken at the same position in the brain stacks. Finally, compare the obtained values to observe any expression changes.

